# Degeneration-Inspired Architectural States Defined by Voronoi Point Spacing and Surface-Mediated Rescue of Osteogenic Dysfunction in 3D-Printed Scaffolds

**DOI:** 10.64898/2026.05.16.725650

**Authors:** Jasmine Carpenter, Pratheesh Kanakarajan Vijaya Kumari, Christopher J Panebianco, Joel D Boerckel, Derrick Dean, Vineeth M Vijayan

## Abstract

Osteoporotic bone degeneration involves progressive deterioration of trabecular microarchitecture, yet most scaffold-based bone tissue engineering studies evaluate osteogenesis in structurally favorable architectures that poorly represent compromised bone environments. Here, we establish a degeneration-inspired Voronoi scaffold platform in which point spacing serves as a single tunable architectural parameter to model transitions from dense mechanically integrated to severely deteriorated trabecular-like microenvironments. Increasing point spacing from 1.25 to 2.5 mm progressively reduced scaffold connectivity and stiffness while shifting deformation behavior from distributed load transfer to localized stress concentration, as confirmed by finite element analysis and mechanical testing. Benchmarking against clinically reported HR-pQCT datasets from postmenopausal women demonstrated that the intermediate 1.75 mm point spacing scaffold represents a clinically relevant compromised trabecular-like state, whereas the 2.5 mm scaffold represents a more severely deteriorated architectural condition. These architecture-dependent mechanical and structural transitions directly regulated hMSC behavior, where high point spacing scaffolds reduced cytoskeletal organization, stress fiber density, and osteogenic mineralization, establishing an architecture-associated osteogenic dysfunction regime. Polydopamine (PDA) coating progressively enhanced cytoskeletal organization and mineralization within architecturally compromised scaffolds without altering scaffold geometry. To quantitatively assess biointerface-mediated functional recovery, a Mineralization Rescue Percentage (MRP) framework was introduced, demonstrating up to 43% restoration of architecture-associated mineralization loss following PDA coating. Collectively, this work establishes a clinically contextualized degeneration-to-rescue biomaterials framework that shifts current scaffold design paradigms beyond structurally favorable architectures toward systematic investigation and functional rescue of architecture-associated osteogenic dysfunction within compromised bone-like microenvironments.

**Graphical Abstract:** 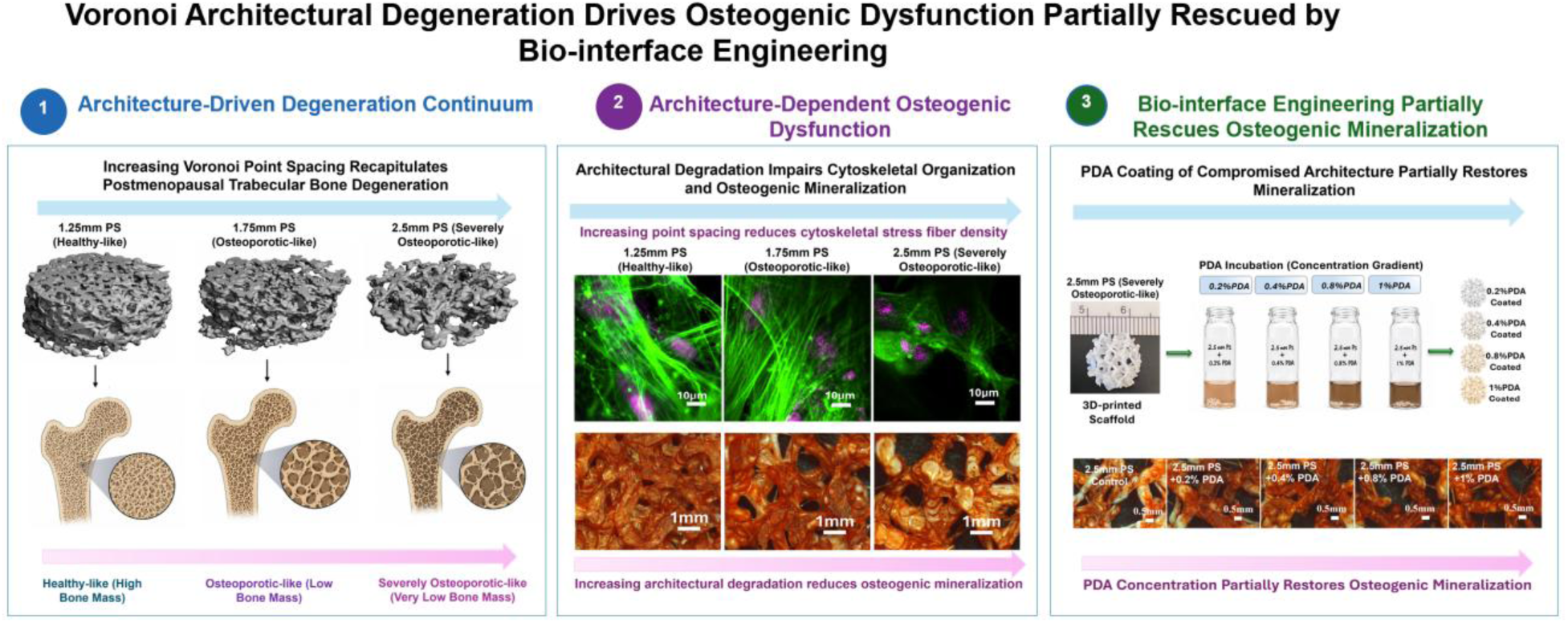

**Statement of Significance:** Most scaffold-based bone tissue engineering studies evaluate osteogenesis in structurally favorable architectures that poorly represent compromised bone microenvironments associated with osteoporosis. Here, a clinically contextualized Voronoi scaffold platform is established in which point spacing serves as a single tunable architectural parameter to model transitions from mechanically integrated to structurally deteriorated trabecular-like states. By decoupling architectural and surface biointerface effects, the study demonstrates that architectural deterioration alone can drive cytoskeletal disruption and osteogenic failure. Importantly, polydopamine-mediated surface engineering partially restored cytoskeletal organization and mineralization within architecturally compromised scaffolds without altering bulk geometry. A Mineralization Rescue Percentage (MRP) framework was further introduced to quantitatively assess biointerface-mediated functional recovery within degeneration-inspired scaffold microenvironments.

## Introduction

Architectural organization is a central determinant of scaffold mechanical competence and cell fate regulation in bone tissue engineering[1, 2]. Beyond bulk material composition, features such as pore size, interconnectivity, and spatial heterogeneity govern load transfer, local stress distributions, and cell–matrix interactions that ultimately influence osteogenic differentiation and matrix mineralization[3–5]. Additive manufacturing has enabled precise control over scaffold architecture, allowing systematic exploration of periodic lattices and stochastic networks designed to approximate trabecular bone[6, 7]. However, most scaffold design strategies achieve “optimization” by simultaneously modifying multiple interdependent parameters such as strut thickness, porosity, unit-cell geometry, gradients, and surface area making it difficult to isolate the specific architectural drivers of mechanical competence and biological function[8, 9].

Voronoi-based lattices have emerged as a particularly attractive class of architected biomaterials due to their inherent stochasticity, tunable connectivity, and resemblance to disordered trabecular architectures[10, 11]. Prior studies have demonstrated the utility of Voronoi scaffolds for achieving bone-relevant transport and mechanical behavior and have combined experimental testing with finite element analysis (FEA) to evaluate mechanical performance and deformation and failure characteristics[12, 13]. Moreover, Voronoi designs have advanced into translational scaffold concepts, including scaffold-guided bone regeneration and topology-driven host-response engineering such as immunomodulation and osteointegration. Despite these advances, existing work commonly evaluates a limited number of discrete design configurations, relies on relative density or porosity as the primary design variable, or treats Voronoi architecture as a static geometry rather than a continuously tunable design space[14–16]. As a result, a key gap remains in establishing a minimal, single-parameter architectural framework that can systematically encode transitions in connectivity, deformation mode, and functional osteogenic performance within a unified material and manufacturing system.

In native bone, gradual architectural deterioration from dense, well-connected trabecular networks to sparse, mechanically compromised structures is associated with marked reductions in load-bearing capacity and increased fracture risk in osteoporosis[17]. Importantly, these transitions occur across a continuum rather than as discrete states, where progressive loss of connectivity and structural redundancy limits the ability of cells to establish mechanically integrated microenvironments necessary for osteogenesis. While numerous scaffold systems have attempted to replicate aspects of trabecular architecture[18, 19], few have been able to systematically model this continuum of architectural degeneration in a controlled and quantifiable manner[20]. Even fewer studies define the point at which architectural degradation leads to functional impairment or osteogenic failure[21].

Despite significant advances in scaffold design and bioactive material development, a critical limitation in current in vitro models is the inability to decouple the relative contributions of architectural mechanics and cell–material interface signaling in governing osteogenic outcomes. In most engineered systems, modifications to scaffold architecture are inherently accompanied by changes in surface chemistry, stiffness, or ligand presentation[22, 23], making it difficult to determine whether observed biological responses arise from structural cues or interfacial bioactivity. This limitation is particularly relevant in the context of bone degeneration, where diseases such as osteoporosis are characterized by progressive architectural deterioration including reduced trabecular connectivity and mechanical competence while the cellular machinery for bone formation remains present but functionally impaired[24, 25].

In this context, there is a critical need for *in vitro* platforms that can independently control architectural and interfacial parameters to determine whether osteogenic dysfunction arises from mechanical disintegration or insufficient cell–material interactions. Such decoupled systems would enable a clearer understanding of structure–function relationships in bone and provide a framework to evaluate whether structurally compromised environments can be functionally modulated through targeted biointerface engineering.

Here, we address this gap by treating Voronoi point spacing as a single continuous architectural design variable governing the transition from mechanically integrated, osteogenically competent microenvironments to structurally compromised architectures with reduced mineralization. By isolating point spacing as the primary geometric parameter, we establish a direct relationship between stochastic architecture, deformation behavior, cytoskeletal organization, and osteogenic function within a unified fabrication framework, enabling identification of architecture-associated functional thresholds beyond which mechanosensitive osteogenic responses become impaired. Building on this framework, we further investigate whether such architecture-induced functional limitations can be partially rescued through biointerface engineering using polydopamine (PDA), a bioinspired surface modification designed to enhance cell–material interactions without altering bulk scaffold geometry [26,27]. Unlike prior studies that primarily evaluate PDA in structurally favorable systems, this work examines whether nanoscale biointerface engineering can restore osteogenic functionality within architecturally compromised microenvironments where mechanobiological responsiveness is diminished.

## 2. Materials and Methods

### 2.1 Designing and printing of Voronoi lattice scaffolds

Irregular stochastic Voronoi lattice scaffolds with varying point spacings (1.25, 1.5, 1.75, 2.0, 2.25, and 2.5 mm) were designed using nTopology software (nTopology Inc., New York, NY, USA). The Voronoi structures were generated using a tolerance of 0.3 mm, an edge length of 0.2 mm, and a growth rate of 2 mm to systematically control architectural features. The point spacing parameter varied from 1.25 to 2.5 mm to generate the six distinct scaffold architectures. Cylindrical scaffolds with a diameter of 14 mm and height of 5 mm were used as the base geometry. The finalized designs were exported as STL files and processed using FlashPrint 5 software (FlashForge, Zhejiang, China) for slicing. Slicing parameters, including layer height and resolution, were optimized to ensure accurate reproduction of the lattice architecture. Scaffolds were fabricated using polylactic acid (PLA) filament (1.75 mm diameter) via fused deposition modeling (FDM) using a Creator Max 3D printer (FlashForge, Zhejiang, China). Printing was performed at a nozzle temperature of 200 °C and a build plate temperature of 50 °C, with a print speed of 30 mm/s and a travel speed of 60 mm/s. These parameters were selected to ensure consistent print fidelity, dimensional accuracy, and structural integrity across all scaffold designs.

### 2.2. Finite element analysis (FEA) of Voronoi lattice scaffolds

Finite element analysis (FEA) was performed to evaluate the mechanical behavior of the different Voronoi scaffolds with varying point spacing under compressive loading. The analysis was conducted on Voronoi structures with varying point spacing, such as 1.25, 1.5, 1.75, 2, 2.25, and 2.5mm. The scaffolds were modeled as isotropic linear elastic materials with material properties corresponding to PLA, including Young’s modulus of 3.5 GPa and Poisson’s ratio of 0.36. A static compression analysis was performed with displacement restraint and force as boundary conditions. A compressive force vector of (0-5000) N was applied to simulate physiological loading conditions. A finite element mesh was generated with appropriate mesh density, and the entity of nodes was defined for the analysis. The simulation evaluated key mechanical parameters, including stress distribution, strain, and displacement within the scaffold structures. This analysis enabled the comparison of mechanical performance across different point spacings to optimize scaffold design.

### 2.3. Microcomputed Tomography (MicroCT) of TPMS Gyroid Scaffolds

High resolution three-dimensional microcomputed tomography (MicroCT) images of the TPMS gyroids were captured in the air with the MicroCT 45 system (SCANCO Medical Ag, Brüttisellen, Switzerland) The imaging parameters applied were: isotropic voxel size of 20 μm, integration time of 300 ms, X-ray intensity of 145 μA, and peak tube voltage of 55 kVp. Visual inspection was performed to accurately identify the threshold used for trabecular morphometry analysis. For noise suppression a three-dimensional gaussian filter of 3.7 with filter support of 4 was used. The volume fraction, strut and pore parameters were quantified using the manufacturer provided software. Parameters are indicated by standard nomenclature for trabecular bone analysis, but applies to scaffold properties (e.g., “bone volume fraction, BV/TV” refers to scaffold volume fraction).

### 2.4 Fluorescence Imaging of Cytoskeletal Organization in hMSCs on Voronoi Scaffolds

To visualize the cytoskeletal organization and cell morphology an actin staining was performed on hMSCs seeded scaffolds. The hMSCs purchased from ATCC (ATCC, USA: Cat no: PCS-500-012) were seeded on Voronoi scaffolds of three different point spacings (1.25mm, 1.75mm, and 2.5mm), as well as the PDA coated scaffolds at a seeding density of 0.7×10^5^ cells. for 48 hours in a CO_2_ incubator (Thermo Scientific, Thermo Fisher Scientific, USA: Model: Forma Series). The scaffolds were washed gently with 1× Phosphate-Buffered Saline (PBS) (Gibco, Thermo Fisher Scientific, USA: Cat no: 14190-144) in a shaker for three times (3 minutes each) to remove culture media. The cells were fixed with 4% paraformaldehyde (Thermo Fisher Scientific, USA: Cat no: J19943-K2) (PFA) for 45 minutes at room temperature. The scaffolds were washed with 1x PBS three times and permeabilized with 0.1% Triton X-100 (Fisher Scientific, USA: Cat no: BP151-500) in PBS for 15 minutes at room temperature. The cells were then incubated with Alexa Fluor 488 Phalloidin in 1% Bovine serum albumin (Thermo Scientific, Thermo Fisher Scientific, USA: Cat no: 37520) (BSA) diluted in PBS at a ratio of 1:400 for 1.5 hours at room temperature to stain the F-actin filaments. Nuclear staining was performed using DRAQ5 (Invitrogen, Thermo Fisher Scientific, USA: Cat no: 65088092) at a dilution of 1:2000 (10μM in PBS) for 15 minutes in the dark. The scaffolds were washed thoroughly with 1xPBS to remove unbound dye, and confocal images were captured using Nikon confocal laser scanning microscope (Nikon Eclipse Ti2). High resolution images were acquired at 4× and 20× magnifications to visualize cell spreading, morphology, actin intensity and stress fiber density on the different scaffold.

### 2.5. Quantitative Analysis of Actin Intensity and Stress Fiber Density

The cytoskeletal organization was assessed on fluorescence images of F-actin stained with phalloidin and imaged using confocal microscopy (Nikon Eclipse Ti2) under identical acquisition settings on all samples. Image analysis was performed using Fiji (ImageJ). Briefly, images were converted to 8-bit grayscale and fixed-size regions of interest (ROIs; 200×200 pixels per side) were applied in cell-covered regions, excluding edges, pores and acellular areas. Actin fluorescence intensity was quantified as the mean gray value after the background subtraction. Stress fiber density was quantified by thresholding the actin channel using a consistently across all samples by calculating the area fraction of filamentous structures within each ROI. Data are presented as mean ± standard error mean (SEM).

### 2.6 Polydopamine coating on 2.5mm Point spacing Voronoi scaffolds

Polydopamine (PDA) coating was performed on 3D-printed Voronoi scaffolds with 2.5 mm point spacing following surface activation. Briefly, scaffolds were subjected to air plasma treatment (Harrick Plasma, PDC-001-HP) to enhance surface wettability. Plasma exposure was carried out at 13.56 MHz RF power (45 W) with a gas flow rate of 40 sccm for 10 min. Following activation, scaffolds were immersed in dopamine hydrochloride solutions at concentrations of 0.2%, 0.4%, 0.8%, and 1% (w/v) (Thermo Fisher Scientific, Waltham, MA, USA; Cat. No. A11136.14) and incubated at 37 °C for 18 h to allow oxidative polymerization and deposition of PDA onto the scaffold surface[27]. After coating, scaffolds were thoroughly rinsed with Milli-Q water (MilliporeSigma, USA) and subsequently sonicated in a bath sonicator (Branson 2510, Danbury, CT, USA) for 5 min to remove loosely bound residues.

### 2.7. Scanning electron microscopy (SEM)

The surface architecture of the scaffolds was examined using a scanning electron microscope (Phenom-XL). The scaffolds were sputter coated with 10nm gold. SEM images were then captured using the backscattered electron detection method. Images were acquired at 5,000x magnification to visualize the surface topography and polydopamine coating.

### 2.8. Alizarin red staining and quantification of mineralization

Alizarin red staining was performed on the Voronoi scaffolds on 7, 14 and 21 days to evaluate the mineralization potential. The hMSCs were seeded on the scaffolds at a seeding density of 0.7×10^5^ cells per scaffold. Cells were incubated in mesenchymal stem cell complete media (Basal media: ATCC, USA: Cat no: PCS-500-030 and growth kit: ATCC, USA: Cat no: PCS-500-041) for five days. Subsequently the stem cell media was removed and washed gently with 1x PBS and supplemented with complete osteogenic differentiation media (StemPro™ Osteogenesis Differentiation Kit: Gibco, Thermo Fisher Scientific, USA: Cat no: A1007201). The osteogenic media change was performed every three days. At specific time points of 7, 14 and 21 day the scaffolds were gently washed with 1x PBS for three times to remove residual medium. The cells were fixed with 4% PFA for 45 minutes at room temperature. Following a 1x PBS wash the scaffolds were incubated with 2% alizarin red s solution (Millipore Sigma, USA: Cat no: TMS008C) for 45 minutes with gentle shaking at room temperature. The unbound dye was carefully washed with 1x PBS. The scaffolds were allowed to dry overnight at room temperature. Stereo microscopic images at 5x magnification were taken using Fisher Scientific stereomicroscope equipped with Motic Cam 2.0. The osteogenic mineralization was quantified using 10% cetylpyridinium chloride (CPC) (MP Biomedicals, USA: Cat no: MP219017780) in 10mM sodium phosphate buffer (pH:7) filtered using 0.2µm syringe filter. 50μl elute was transferred to a 96 well plate afterincubation for 15 minutes with 10% CPC. Optical density was read at 560nm in BioTek Cytation 3 (BioTek, USA) imaging reader. DNA quantification for normalization was performed using a PicoGreen assay on duplicate scaffolds seeded concurrently at the same seeding density and under identical culture conditions. The alizarin red absorbance values were normalized to the corresponding DNA content and data are presented as mean ± standard error mean (SEM).

### 2.9 Quantification of Mineralization Rescue Percentage (MRP)

To quantitatively assess the extent to which surface modification influences mineralization in structurally compromised scaffolds, a Mineralization Rescue Percentage (MRP) metric was defined based on Alizarin Red S (ARS) quantification normalized to DNA content. Mineralization values were first normalized using a PicoGreen assay to account for differences in cell number across samples. Briefly, ARS values were divided by the corresponding DNA content measured via PicoGreen fluorescence, yielding DNA-normalized mineralization values for each condition.

MRP was then calculated by normalizing the mineralization of treated scaffolds to the difference between a dense (high-connectivity) reference scaffold and a structurally compromised scaffold. The metric is defined as:

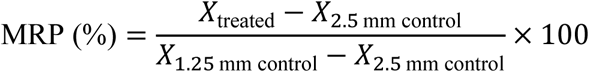

where:

- *X*_1.25 mm control_ represents the DNA-normalized ARS value of the dense scaffold (1.25 mm point spacing), used as the reference healthy bone condition,
- *X*_2.5 mm control_ represents the DNA-normalized ARS value of the structurally compromised scaffold (2.5 mm point spacing, untreated), and
- *X*_treated_ corresponds to the DNA-normalized ARS value of 2.5 mm scaffolds after different concentrations of PDA surface modification (0.2,0.4,0.8 and 1%).

All values were averaged across replicates before MRP calculation. The denominator (*X*_1.25_−*X*_2.5_)represents the reduction in mineralization associated with architectural differences, and the MRP expresses the fraction of this reduction that is recovered following treatment.

An MRP value of 0% indicates no change relative to the compromised scaffold, whereas a value of 100% corresponds to mineralization levels comparable to the dense reference condition. Values between 0% and 100% represent partial recovery. The MRP is used here as a relative metric to compare treatment-induced changes in mineralization within the context of architecture-driven baseline differences, rather than as an absolute measure of osteogenic potential.

## 3. Results

### 3.1 Voronoi point spacing defines a continuous architectural design space

The heterogeneous and irregular nature of native trabecular bone motivated the use of a Voronoi tessellation strategy to design three-dimensional scaffolds with controlled architectural heterogeneity [28]. To systematically capture this architectural variation, a scalar field–based Voronoi approach was employed in which point spacing was defined as the sole architectural design parameter and varied continuously from 1.25 to 2.5 mm (Figure 1a). This strategy enabled continuous modulation of scaffold architecture while preserving identical material composition and fabrication conditions.

**Figure 1.**
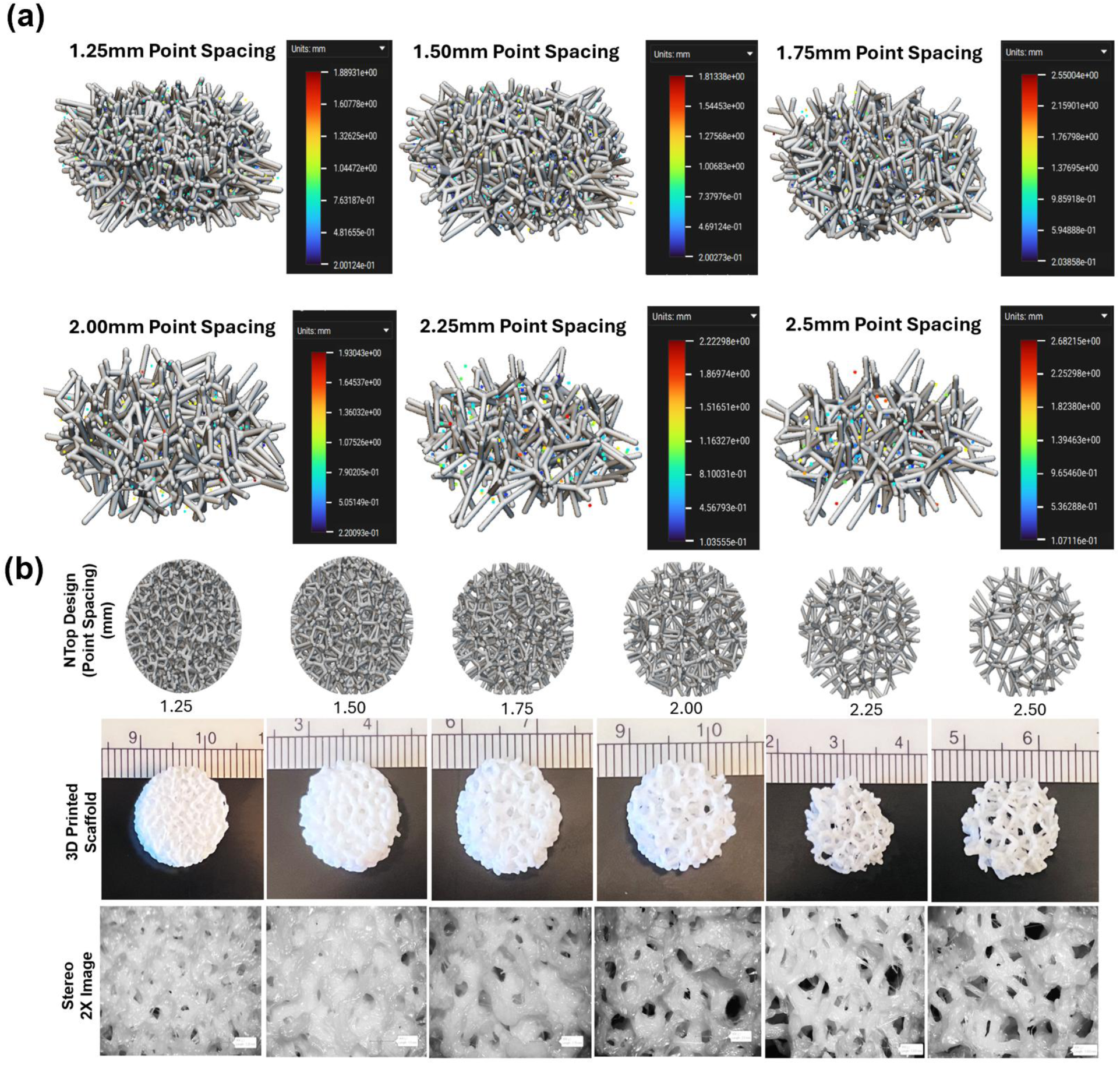
*(a)* Scalar field mapping illustrating Voronoi tessellations generated using progressively increasing point spacing (1.25–2.5 mm), with color maps indicating local pore size distributions and the corresponding three-dimensional Voronoi lattice architectures derived from each tessellation. *(b)* nTopology-rendered Voronoi scaffold designs (top), representative photographs of 3D-printed polylactic acid (PLA) scaffolds (middle), and stereomicroscopy images (bottom) confirming the faithful translation of digitally programmed architectures into physical constructs. Increasing point spacing transitions the scaffold morphology from dense, highly connected networks to more open, porous architectures with reduced connectivity and larger characteristic pore sizes.

As shown in Figure 1a, increasing point spacing systematically alters the spatial distribution of Voronoi seed points, resulting in corresponding changes in local pore size distribution and network topology. These digitally generated architectures were translated into three-dimensional lattice structures, revealing a progressive transition from dense, highly interconnected networks at low point spacing to increasingly sparse and heterogeneous architectures at higher point spacing.

The fidelity of this design-to-fabrication process was confirmed through 3D printing of PLA scaffolds, where the printed constructs closely matched the computational designs (Figure 1b). Macroscopic imaging and stereomicroscopy further demonstrated that increasing point spacing produces larger pore openings, reduced node density, and decreased structural connectivity. Collectively, these results establish Voronoi point spacing as a robust and scalable single-parameter design handle for generating a continuum of architecturally distinct scaffold microenvironments.

### 3.2 Point spacing governs deformation mode and mechanical competence and enables selection of representative structural states

To determine how architectural modulation influences mechanical behavior, combined experimental and finite element analyses were performed across the full range of point spacings.

Finite element simulations demonstrated progressive changes in deformation behavior with increasing point spacing (Figure 2a). Scaffolds with low point spacing (1.25 mm) exhibited relatively uniform displacement distributions under uniaxial compression, consistent with more distributed load transfer across a connected architectural network. In contrast, higher point spacing scaffolds showed increasingly localized deformation patterns, with displacement concentrated within fewer structural elements. Corresponding von Mises stress distributions also became progressively more heterogeneous with increasing point spacing, indicating reduced load-sharing capacity and increased stress localization within sparsely connected architectures. Experimental compression testing demonstrated trends consistent with the FEM simulations. Experimental Young’s modulus progressively decreased with increasing point spacing, from approximately 1.76 GPa at 1.25 mm PS to 0.30 GPa at 2.5 mm PS (Figure S1a). FE-predicted modulus values followed a similar monotonic reduction across the architectural range (Figure S1b), supporting the ability of the computational model to capture the overall mechanical trends associated with increasing structural sparsity. Quantitative FEM analysis further supported these observations (Figure S2A–C). Mean displacement increased from 0.18 mm at 1.25 mm PS to 1.36 mm at 2.5 mm PS, while the maximum-to-mean stress ratio increased from 9.26 to 19.56, indicating progressively concentrated stress distributions at larger point spacings. Mean von Mises stress also increased with increasing point spacing, accompanied by higher stress variability, as reflected by larger standard deviation values.

**Figure 2.**
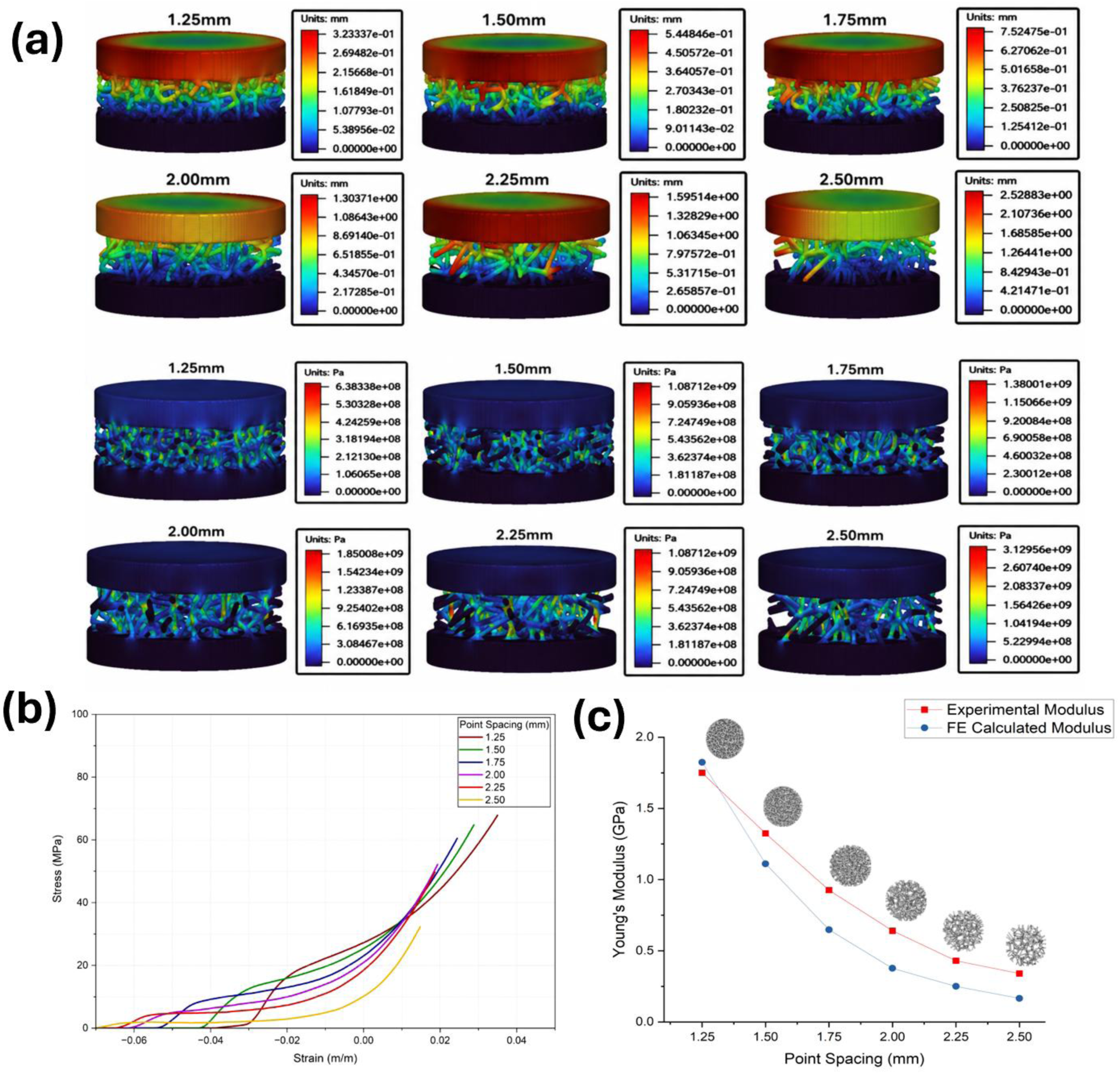
(a) Finite element analysis of Voronoi scaffolds under uniaxial compression showing displacement distributions (top; mm) and von Mises stress distributions (bottom; Pa) for point spacings ranging from 1.25 to 2.50 mm. (b) Representative experimental compressive stress–strain curves for each point spacing condition. (c) Comparison of experimentally measured and finite element–calculated Young’s modulus as a function of point spacing, showing a progressive reduction in scaffold stiffness with increasing point spacing.

Stress–strain curves further demonstrated a progressive reduction in scaffold stiffness with increasing point spacing (Figure 2b). Comparison of experimental and FE-calculated modulus values showed agreement in the overall mechanical response trend across the design space (Figure 2c). Together, these findings demonstrate that increasing Voronoi point spacing progressively shifts scaffold behavior from mechanically integrated toward structurally compromised states characterized by reduced stiffness and localized stress concentration.

**Figure 3:**
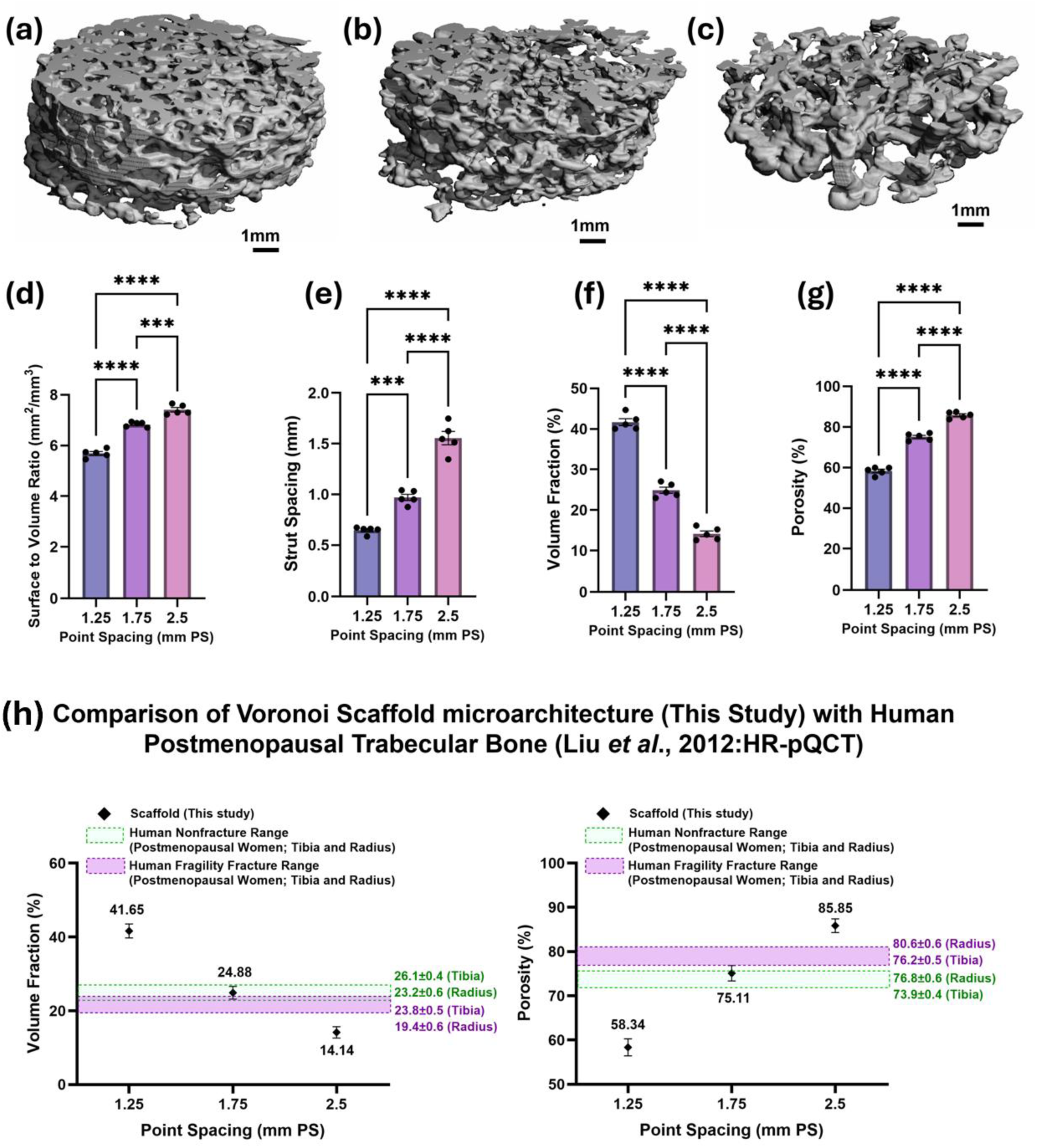
Architectural characterization and clinical benchmarking of Voronoi scaffolds with varying point spacing (PS). (a–c) Representative three-dimensional μCT reconstructions of Voronoi scaffolds with PS values of 1.25, 1.75, and 2.5 mm, respectively. Quantitative analysis of (d) surface area-to-volume ratio (BS/BV-equivalent), (e) scaffold strut spacing (Tb.Sp-equivalent), (f) scaffold volume fraction (BV/TV-equivalent), and (g) porosity demonstrating progressive architectural transition with increasing PS. (h) Comparison of scaffold-derived architectural parameters with clinically reported HR-pQCT trabecular bone microarchitectural data from postmenopausal women with and without fragility fractures reported by Liu et al [29]. The intermediate 1.75 mm PS scaffold exhibited scaffold volume fraction and porosity values within the clinically reported postmenopausal trabecular bone microarchitectural range at the distal tibia and radius, whereas the 1.25 mm and 2.5 mm PS scaffolds represented denser and severely deteriorated architectural conditions, respectively. Scaffold-derived parameters are presented as architectural analogs of clinically reported bone morphometry metrics, including BS/BV, Tb.Sp, and BV/TV. Data are presented as mean ± SD (*p < 0.05, **p < 0.01, ***p < 0.001, ****p < 0.0001).

Based on the combined architectural, mechanical, and FEM analyses, 1.25 mm, 1.75 mm, and 2.5 mm PS scaffolds were selected as representative dense, intermediate, and highly compromised architectural states, respectively, for subsequent biological investigations. To demonstrate the practical scalability of this platform, scaffolds representing these selected point spacings (1.25, 1.75, and 2.5 mm) were further integrated into standard multiwell plate formats (24-, 48-, and 96-well plates) (Figure S3). The ability to accommodate multiple scaffold conditions within commonly used culture formats highlights the compatibility of this system with high-throughput experimental workflows and supports its use as a platform for systematic evaluation of structure–function relationships.

### 3.3 Architectural Benchmarking of Voronoi Scaffolds Against Clinically Reported Human Trabecular Bone Microarchitecture

Representative three-dimensional μCT reconstructions of Voronoi scaffolds with point spacing (PS) values of 1.25, 1.75, and 2.5 mm are shown in Fig. 4a–c. Progressive increases in PS produced distinct architectural transitions from a dense interconnected trabecular-like morphology toward a sparse and structurally compromised porous network. Quantitative μCT analysis demonstrated that increasing PS significantly increased the surface area-to-volume ratio (BS/BV-equivalent) and scaffold strut spacing (Tb.Sp-equivalent) of the scaffolds (Fig. 4d,e). Specifically, the BS/BV-equivalent increased from 5.69 ± 0.17 mm²/mm³ in the 1.25 mm PS group to 7.41 ± 0.19 mm²/mm³ in the 2.5 mm PS group, while scaffold strut spacing (Tb.Sp-equivalent) increased from 0.65 ± 0.03 mm to 1.56 ± 0.15 mm.

**Figure 4:**
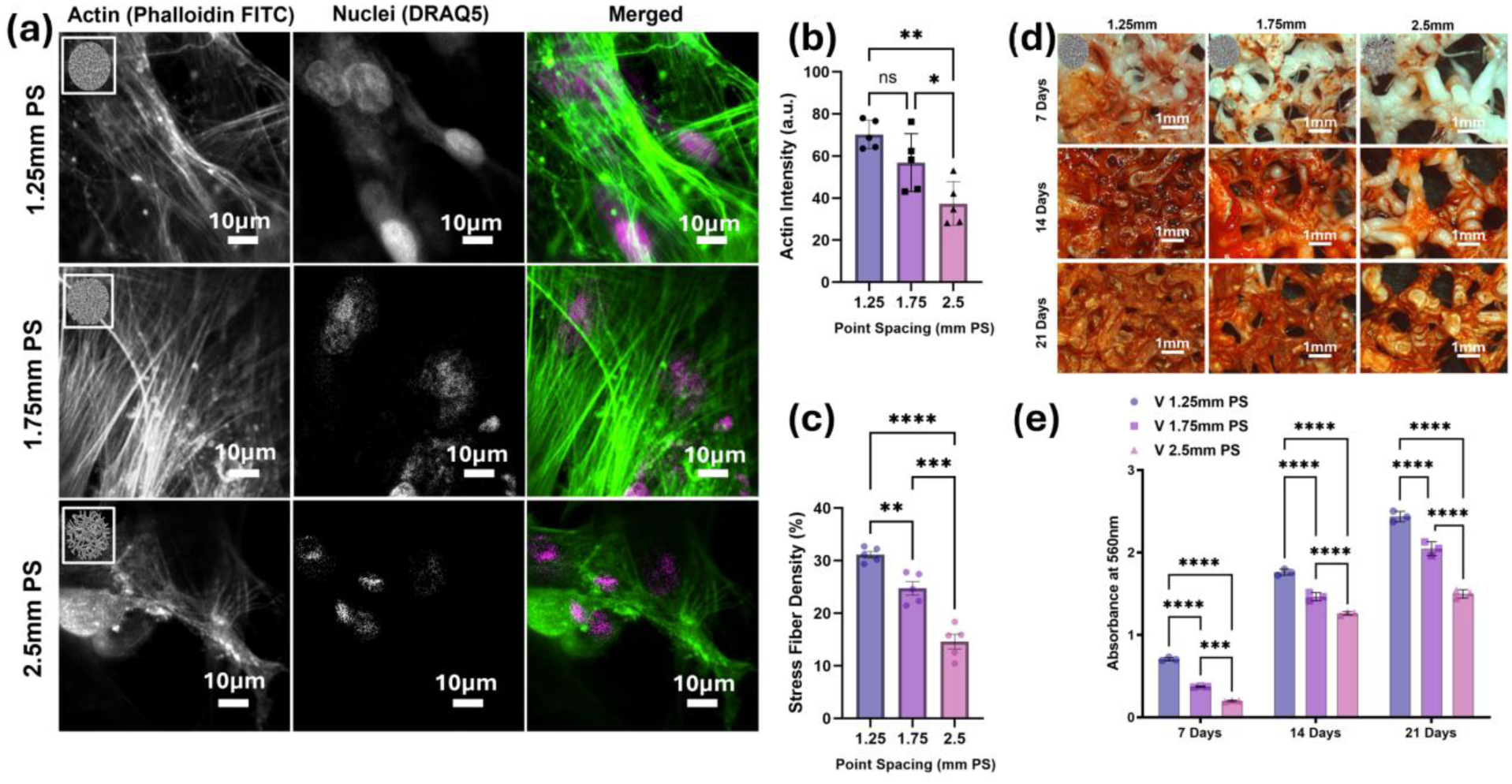
Architecture-dependent cytoskeletal organization and osteogenic mineralization in Voronoi scaffolds with varying point spacing (PS). (a) Representative confocal fluorescence images of hMSCs cultured on 1.25, 1.75, and 2.5 mm PS scaffolds showing F-actin organization (Phalloidin-FITC), nuclei (DRAQ5), and merged images. Lower PS scaffolds exhibited greater cell spreading and more developed actin stress fiber organization, whereas the 2.5 mm PS scaffold showed reduced cytoskeletal organization. Scale bars = 10 μm. Quantification of (b) actin intensity and (c) stress fiber density. (d) Representative stereomicroscopy images of Alizarin Red S staining at 7, 14, and 21 days demonstrating reduced mineralization with increasing PS. Scale bars = 1 mm. (e) Quantification of normalized mineralization absorbance at 560 nm showing time-dependent osteogenic mineralization across scaffold groups. Data are presented as mean ± SD (*p < 0.05, **p < 0.01, ***p < 0.001, ****p < 0.0001).

In contrast, scaffold volume fraction (BV/TV-equivalent) progressively decreased with increasing PS, accompanied by a corresponding increase in porosity (1 − BV/TV-equivalent) (Fig. 4f,g). The 1.25 mm PS scaffold exhibited the highest scaffold volume fraction (41.66 ± 1.94%) and lowest porosity (58.34 ± 1.94%), whereas the 2.5 mm PS scaffold demonstrated markedly reduced scaffold volume fraction (14.15 ± 1.55%) and elevated porosity (85.85 ± 1.55%). The intermediate 1.75 mm PS scaffold exhibited a scaffold volume fraction of 24.88 ± 1.73% and porosity of 75.12 ± 1.73%, representing an architecturally intermediate state between the dense and highly deteriorated scaffold conditions.

To evaluate the physiological relevance of these engineered architectures, scaffold-derived microarchitectural parameters obtained from high-resolution microcomputed tomography (μCT) analysis were benchmarked against clinically reported high-resolution peripheral quantitative computed tomography (HR-pQCT) datasets from postmenopausal women with and without fragility fractures reported by Liu et al. [29] (Fig. 4h). Importantly, scaffold volume fraction, scaffold strut spacing, and BS/BV-equivalent represent the corresponding architectural analogs of clinically reported bone morphometry parameters BV/TV, Tb.Sp, and BS/BV, respectively. Comparison of scaffold volume fraction and porosity demonstrated that the intermediate 1.75 mm PS scaffold fell within the clinically reported postmenopausal trabecular bone microarchitectural range at the distal tibia and radius, particularly near the overlap region between fracture and non-fracture cohorts. In contrast, the 1.25 mm PS scaffold represented a substantially denser trabecular-like architecture with higher scaffold volume fraction and lower porosity than the reported clinical ranges, whereas the 2.5 mm PS scaffold represented a more severely deteriorated architectural condition characterized by excessive porosity and markedly reduced scaffold volume fraction beyond the average values reported for the fragility fracture cohort.

Collectively, these findings demonstrate that controlled modulation of Voronoi PS enables reproducible tuning of scaffold microarchitecture across dense, clinically relevant compromised, and severely deteriorated trabecular-like architectural states associated with degenerative bone conditions.

### 3.4 Architecture-Dependent Cytoskeletal Organization and Osteogenic Mineralization in Voronoi Scaffolds

To evaluate how architecture-dependent scaffold microenvironments influence early cytoskeletal organization, hMSCs cultured on Voronoi scaffolds with varying point spacing were analyzed using confocal fluorescence imaging (Figure 4a). Scaffolds with low (1.25 mm) and intermediate (1.75 mm) point spacing exhibited extensive cell spreading together with well-developed and interconnected F-actin networks. In contrast, the high point spacing scaffold (2.5 mm) displayed reduced cytoskeletal continuity, with cells exhibiting more restricted attachment and simplified actin organization. Quantitative analysis further demonstrated that both actin intensity and stress fiber density progressively decreased with increasing point spacing (Figure 4b,c), indicating that reduced structural connectivity and larger pore gaps negatively influence cytoskeletal organization within the scaffold microenvironment.

To further assess whether these architecture-dependent differences influenced osteogenic function, mineralization was evaluated using Alizarin Red S staining over 7, 14, and 21 days (Figure 4d). Scaffolds with lower point spacing exhibited greater and more spatially distributed mineral deposition across all time points, whereas the 2.5 mm scaffold consistently showed reduced mineralization. Quantitative absorbance analysis confirmed a significant decrease in mineralization with increasing point spacing at each time point (Figure 4e). Collectively, these findings demonstrate that increasing Voronoi point spacing progressively alters scaffold structural organization, resulting in reduced cytoskeletal organization and impaired osteogenic mineralization within architecturally compromised scaffold microenvironments.

### 3.5 Surface biointerface engineering enables partial rescue of osteogenic function

Having established that increased point spacing generates mechanically compromised, osteoporotic-like architectures, we next evaluated a surface-mediated rescue strategy. Polydopamine (PDA) coatings were applied to high point spacing scaffolds (2.5 mm), representing the most structurally compromised condition, to assess whether surface modification can enhance osteogenic mineralization relative to this baseline and in comparison to the low point spacing (1.25 mm) control. Visual inspection during incubation revealed progressive darkening of scaffolds with increasing PDA concentration (0.2–1.0%) after 18 h, consistent with concentration-dependent PDA formation (Figure S4). Stereomicroscopy further confirmed uniform coating development while preserving the underlying lattice geometry.

Representative photographs demonstrated progressive darkening of the scaffolds with increasing PDA concentration following 18 h incubation, consistent with concentration-dependent PDA deposition (Figure 5A,B). Stereomicroscopy further confirmed that PDA coating was achieved without disrupting the overall lattice geometry or structural integrity of the scaffolds. SEM analysis (Figure 5C–G) revealed progressive surface coverage with increasing PDA concentration. Lower concentrations (0.2–0.4%) exhibited relatively discontinuous surface deposition, whereas higher concentrations (0.8–1.0%) produced increasingly uniform and conformal coatings across the scaffold surface. Collectively, these observations demonstrate controlled concentration-dependent PDA functionalization while maintaining the underlying scaffold architecture, thereby enabling investigation of surface-mediated biological modulation independent of architectural variation.

**Figure 5.**
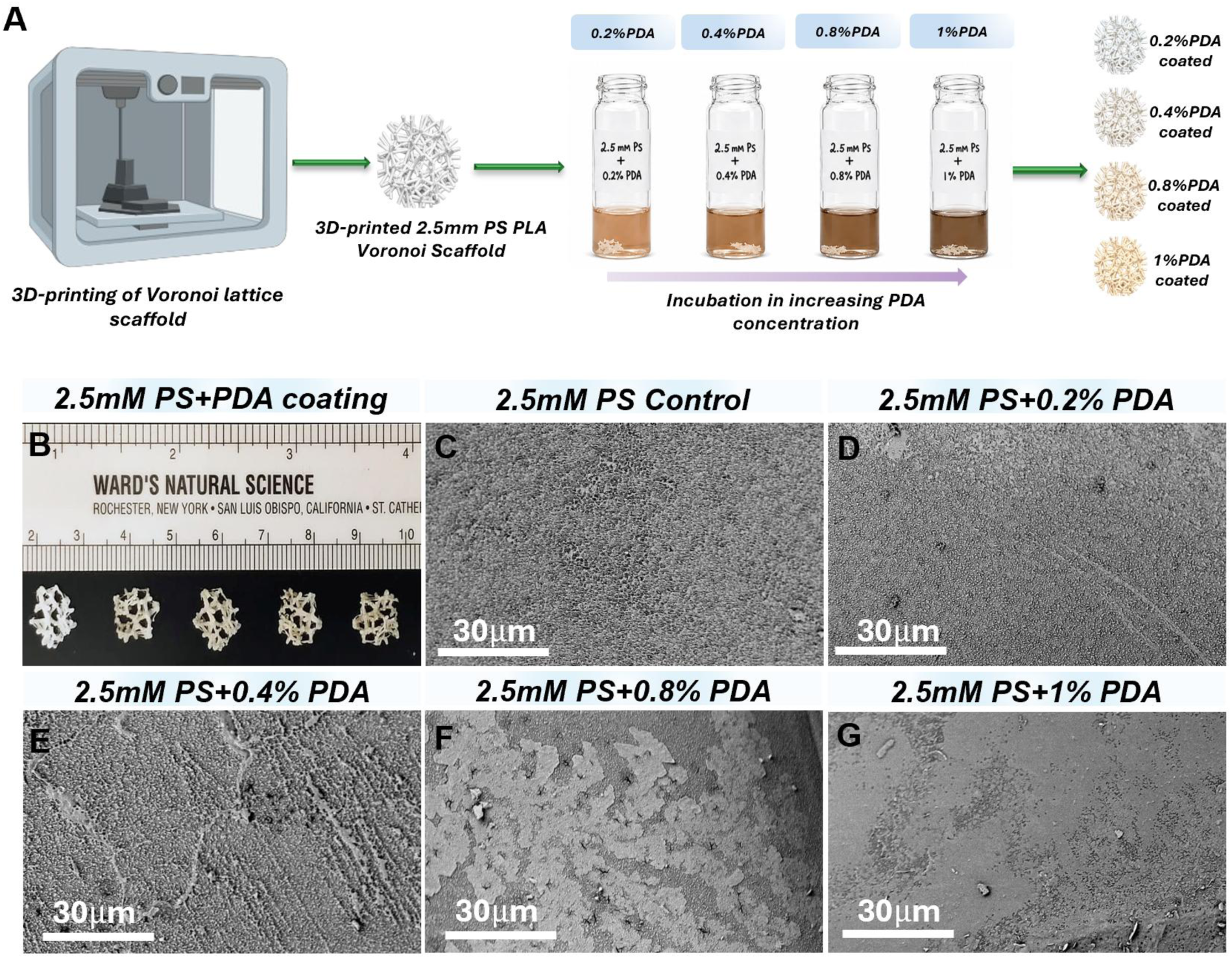
(A) Schematic illustration of the fabrication and surface functionalization process of 3D-printed Voronoi scaffolds with 2.5 mm point spacing (PS), including scaffold printing followed by incubation in increasing polydopamine (PDA) concentrations (0.2%, 0.4%, 0.8%, and 1%) to achieve concentration-dependent surface modification. Representative images of uncoated and PDA-coated scaffolds demonstrating macroscopic appearance following surface treatment (B). (C–G) Scanning electron microscopy (SEM) images of scaffold surfaces showing progressive PDA deposition and increasing surface coverage with increasing PDA concentration. Scale bars = 30 μm.

Three-dimensional confocal reconstructions further demonstrated architecture-dependent differences in hMSC organization across the scaffold groups (Figure 6a). The 1.25 mm PS scaffolds exhibited greater cellular spreading and interconnected actin organization, whereas the 2.5 mm PS control scaffolds showed reduced cellular continuity and more localized cytoskeletal organization. PDA-coated 2.5 mm PS scaffolds demonstrated progressively enhanced cellular spreading and cytoskeletal organization with increasing PDA concentration while maintaining the underlying scaffold architecture. Depth-coded reconstructions further indicated differences in the spatial distribution and organization of cells throughout the scaffold microenvironment (Figure 6b).

**Figure 6:**
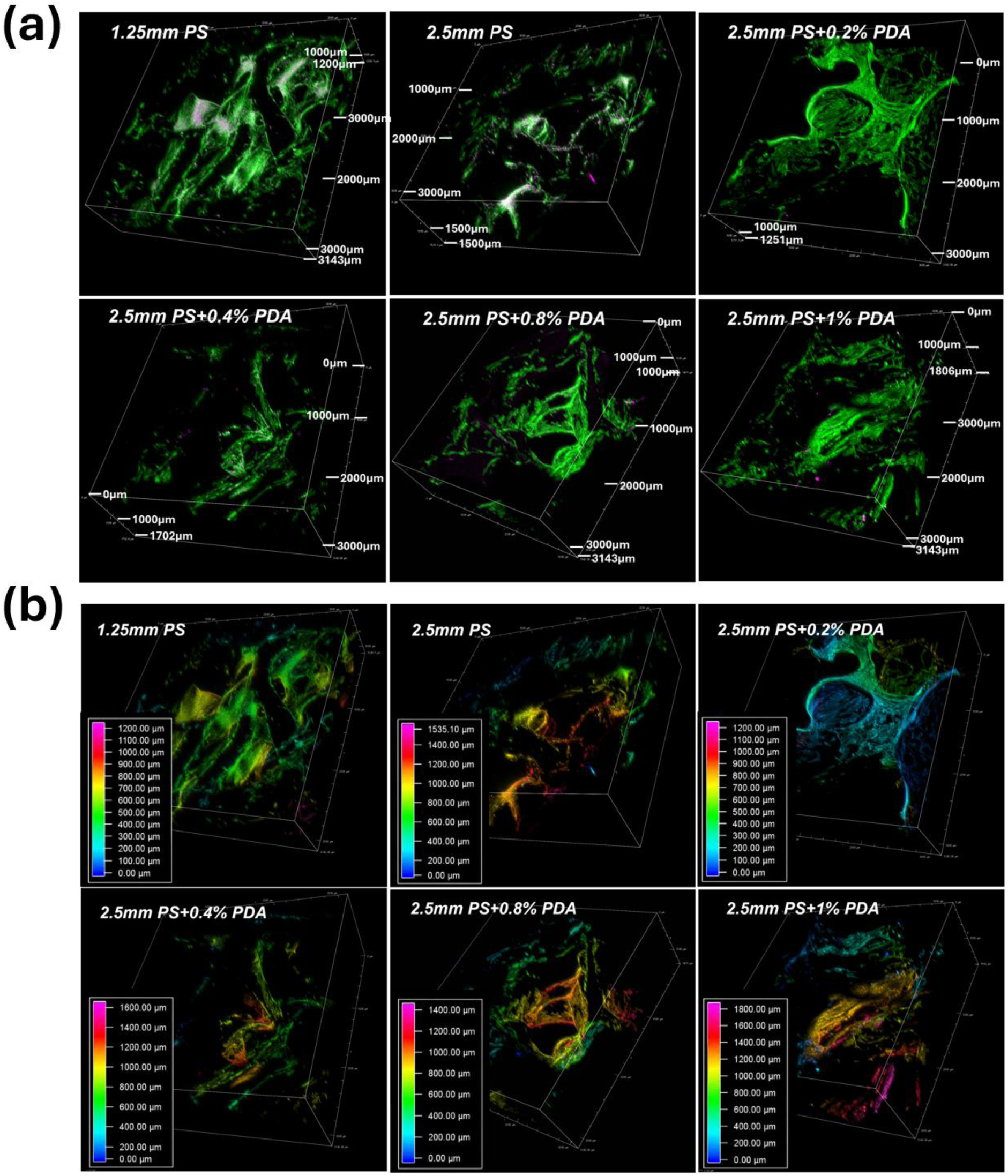
Three-dimensional confocal reconstruction and depth-resolved cytoskeletal organization of hMSCs cultured on degeneration-inspired Voronoi scaffolds with PDA-mediated surface rescue. (a) Representative 3D confocal reconstructions of hMSCs cultured on 1.25 mm PS control scaffolds, 2.5 mm PS control scaffolds, and 2.5 mm PS scaffolds coated with increasing concentrations of polydopamine (PDA; 0.2–1.0%). The 1.25 mm PS scaffold exhibited extensive cell spreading and interconnected actin organization, whereas the 2.5 mm PS control scaffold showed reduced cellular coverage and disrupted cytoskeletal organization. PDA coating progressively enhanced cell spreading and cytoskeletal continuity within the architecturally compromised 2.5 mm PS scaffolds. (b) Depth-coded 3D reconstructions demonstrating spatial organization and scaffold infiltration of hMSCs across the different scaffold conditions. Increased PDA concentration promoted improved cellular distribution and depth-associated cytoskeletal organization within the compromised scaffold microenvironment. Scale bars and depth scales are indicated in the corresponding panels.

Confocal fluorescence imaging demonstrated architecture-dependent differences in hMSC cytoskeletal organization across the scaffold groups (Figure 7a). The 1.25 mm PS scaffolds exhibited greater cell spreading and more developed actin stress fiber organization, whereas the 2.5 mm PS control scaffolds showed reduced cytoskeletal continuity and simplified actin organization. PDA coating of the 2.5 mm PS scaffolds progressively enhanced actin organization with increasing PDA concentration. Quantitative analysis further demonstrated reduced actin intensity and stress fiber density in the untreated 2.5 mm PS scaffolds relative to the 1.25 mm PS condition, while PDA-coated groups exhibited concentration-dependent increases in stress fiber density (Figure 7b,c).

**Figure 7.**
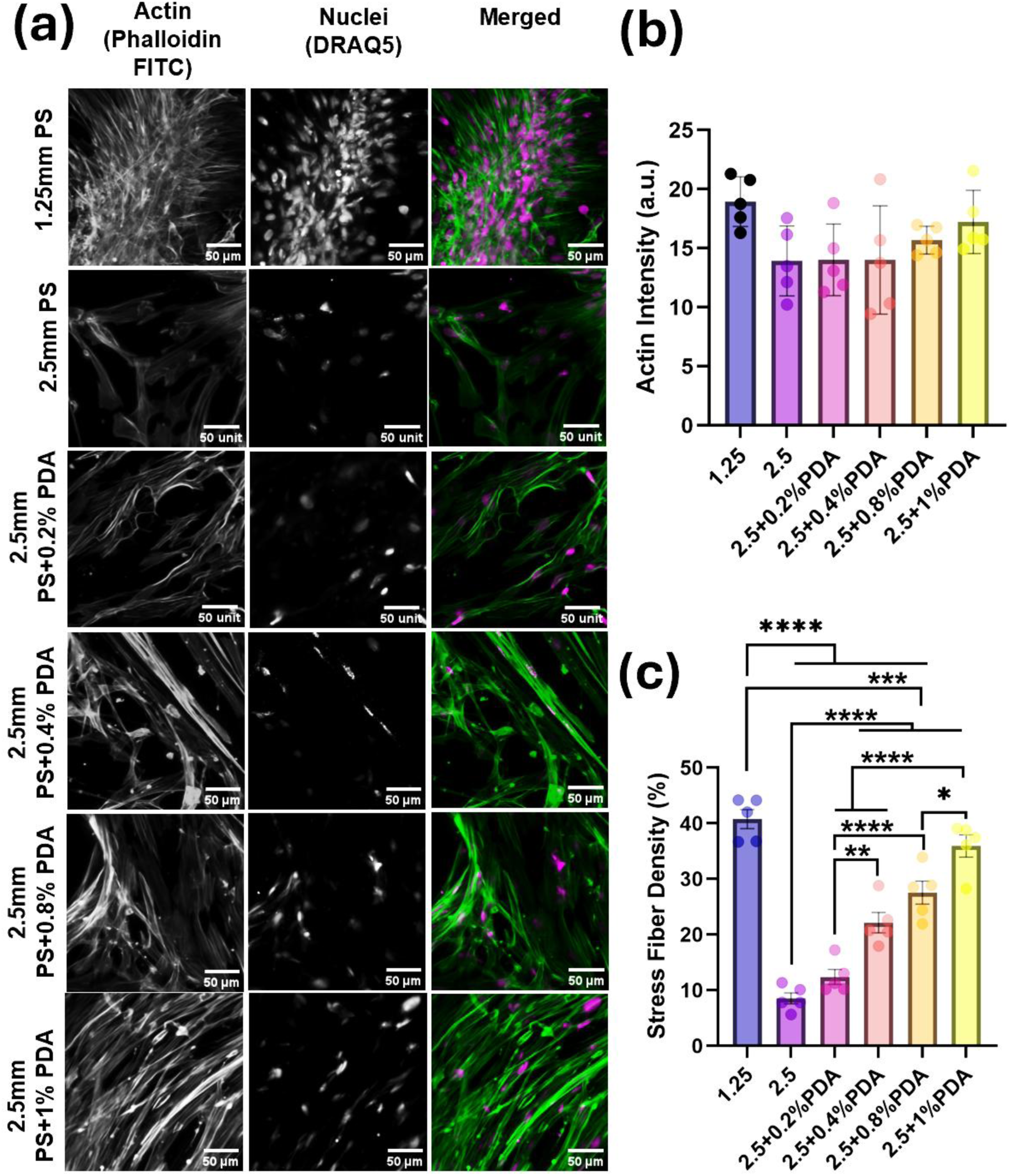
PDA-mediated modulation of cytoskeletal organization in degeneration-inspired Voronoi scaffolds. (a) Representative confocal fluorescence images of hMSCs cultured on 1.25 mm point spacing (PS) scaffolds, untreated 2.5 mm PS scaffolds, and 2.5 mm PS scaffolds coated with increasing concentrations of polydopamine (PDA; 0.2–1.0%), showing F-actin organization (Phalloidin-FITC), nuclei (DRAQ5), and merged images. The 1.25 mm PS scaffolds exhibited greater cytoskeletal organization, whereas untreated 2.5 mm PS scaffolds showed reduced actin continuity. PDA coating progressively improved actin organization within the 2.5 mm PS scaffolds. Scale bars = 50 μm. Quantification of (b) actin intensity and (c) stress fiber density demonstrating concentration-dependent recovery of cytoskeletal organization following PDA coating. Data are presented as mean ± SD (*p < 0.05, **p < 0.01, ***p < 0.001, ****p < 0.0001).

To quantify the extent of mineralization recovery, a Mineralization Rescue Percentage (MRP) metric was defined based on DNA-normalized Alizarin Red absorbance using the 1.25 mm PS scaffold as the reference condition and the untreated 2.5 mm PS scaffold as the reduced-mineralization condition (Figure 8c). Consistent with the qualitative staining observations (Figure 8a), the 2.5 mm PS scaffold exhibited substantially reduced mineralization relative to the 1.25 mm PS scaffold, indicating impaired osteogenic mineral deposition within the architecturally compromised condition (Figure 8b). PDA coating resulted in a concentration-dependent increase in mineralization within the 2.5 mm PS scaffolds, with relatively limited recovery at lower PDA concentrations (0.2% and 0.4%) and progressively greater recovery at higher concentrations (0.8% and 1.0%). The highest PDA concentration (1.0%) exhibited an MRP of approximately 43%, indicating partial restoration of osteogenic mineralization within the compromised scaffold microenvironment. These findings suggest that PDA-mediated surface biointerface engineering can partially improve osteogenic functionality within architecturally compromised scaffold microenvironments without altering the underlying scaffold geometry.

**Figure 8:**
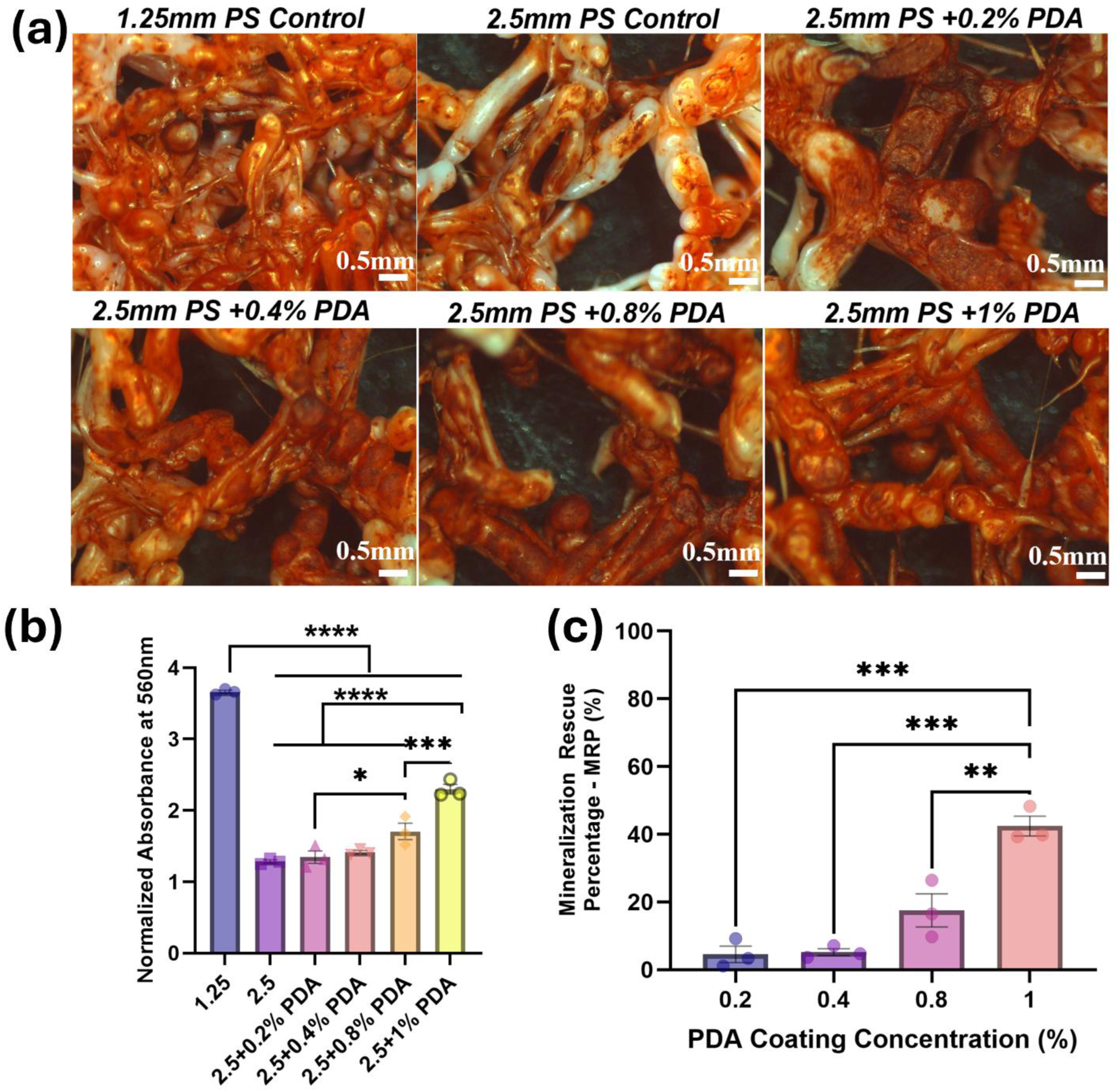
PDA-mediated rescue of osteogenic mineralization in degeneration-inspired Voronoi scaffolds. (a) Representative stereomicroscopy images of Alizarin Red S staining showing osteogenic mineralization in 1.25 mm PS control scaffolds, 2.5 mm PS control scaffolds, and 2.5 mm PS scaffolds coated with increasing concentrations of polydopamine (PDA; 0.2–1.0%). The 2.5 mm PS control scaffold exhibited reduced mineral deposition relative to the 1.25 mm PS scaffold, whereas PDA-coated 2.5 mm PS scaffolds demonstrated progressive recovery of mineralization with increasing PDA concentration. Scale bars = 0.5 mm. (b) Quantification of normalized mineralization absorbance at 560 nm showing concentration-dependent increases in mineralization following PDA coating. (c) Mineralization Rescue Percentage (MRP) analysis demonstrating partial restoration of osteogenic mineralization within architecturally compromised scaffolds following PDA-mediated surface biointerface engineering. Data are presented as mean ± SD (*p < 0.05, **p < 0.01, ***p < 0.001, ****p < 0.0001).

The introduction of the MRP metric provides a framework for comparing treatment-induced changes relative to architecture-driven baseline differences. By expressing mineralization as a fraction of the reduction associated with increased point spacing, this approach enables relative assessment of functional changes across scaffold conditions. While the MRP does not represent an absolute measure of osteogenic potential, it facilitates comparison of surface modification effects within a defined architectural context.

The substantial reduction in mineralization observed in the 2.5 mm scaffold is consistent with the corresponding changes in mechanical behavior associated with increased point spacing. As shown in Figure 2a and b higher point spacing is associated with increased deformation and greater variability in stress distribution, which may influence the local mechanical environment experienced by cells. Within this context, PDA coating is associated with increased mineral deposition in a dose-dependent manner, suggesting that surface chemistry may influence cell–material interactions under these conditions.

However, the observed recovery remains partial even at higher PDA concentrations (∼43% at 1% PDA), indicating that surface modification alone does not fully restore mineralization to levels observed in the denser scaffold. This suggests that architectural features continue to influence the overall mineralization response. Together, these observations support a framework in which scaffold architecture defines the baseline structural and mechanical environment, while surface modification influences cellular responses within that context.

To summarize the relationships identified across structural, cellular, and functional analyses, a schematic overview is presented in Figure 8. Increasing Voronoi point spacing progressively alters scaffold microarchitecture, resulting in reduced structural connectivity and corresponding shifts in μCT-derived parameters. These architectural changes are associated with differences in cytoskeletal organization, where lower point spacing supports more developed actin networks, while higher point spacing is associated with reduced cytoskeletal organization. Consistent with these trends, osteogenic mineralization decreases with increasing point spacing. Surface modification through PDA coating is associated with a partial, concentration-dependent increase in mineralization within the high point spacing condition. Together, this schematic integrates the observed relationships between scaffold architecture, cytoskeletal organization, and osteogenic function, and illustrates how surface modification modulates these responses within a fixed architectural context.

## 4. Discussion

### 4.1 Architectural Control of Mechanical Response and Structural Benchmarking Against Human Degenerative Trabecular Bone Microenvironments

Collectively, Figures 1–3 demonstrate that Voronoi point spacing serves as a primary geometric parameter governing scaffold mechanical behavior and structural organization. Prior studies using Voronoi and other lattice-based scaffold designs have largely focused on recapitulating the heterogeneous, trabecular-like architecture of native cancellous bone, optimizing porosity and mechanical properties, or generating patient-specific bone-mimetic scaffolds with structural similarity to healthy trabecular bone [30–32]. Similarly, TPMS and gyroid lattice scaffolds have been widely investigated because their interconnected porous architectures and smooth curvature can support bone tissue engineering applications [33–35]. However, most of these approaches use architectural complexity to approximate or optimize normal trabecular-like environments rather than to model progressive architectural deterioration associated with compromised bone diseases. In contrast, the present work treats Voronoi point spacing as a continuous and experimentally controllable architectural variable to reproduce degeneration-inspired trabecular-like states relevant to osteoporosis and skeletal fragility. Increasing point spacing produced a progressive reduction in compressive stiffness, accompanied by a transition from distributed stress profiles to increasingly localized stress concentrations. The agreement between experimental and finite element–predicted modulus trends further supports that point spacing captures the dominant architectural determinants of mechanical response across the scaffold designs.

Based on these mechanically defined regimes, three representative conditions (1.25, 1.75, and 2.5 mm) were selected to represent low, intermediate, and high point spacing states. μCT-derived structural characterization demonstrated a progressive transition toward more porous and less connected architectures with increasing point spacing, including increased scaffold strut spacing and porosity together with reduced scaffold volume fraction. These trends are consistent with clinically reported trabecular deterioration in postmenopausal women with fragility fractures [29], where skeletal fragility is associated with impaired trabecular microarchitecture beyond bone mineral density alone. Osteoporosis is characterized by reduced BV/TV, increased trabecular separation (Tb.Sp), and elevated porosity, all of which contribute to skeletal fragility and fracture susceptibility [36,37]. Accordingly, clinically reported HR-pQCT datasets from postmenopausal women were selected as a benchmarking framework because postmenopausal osteoporosis represents one of the most clinically relevant forms of trabecular degeneration associated with fragility fractures.

Liu et al. demonstrated that postmenopausal women with fragility fractures exhibited significantly lower BV/TV and altered trabecular organization compared to non-fracture controls [29]. Consistent with these observations, increasing Voronoi point spacing progressively transformed the scaffold architecture from a dense interconnected network toward a sparse and structurally compromised morphology characterized by reduced scaffold volume fraction and increased scaffold strut spacing. Specifically, increasing point spacing from 1.25 mm to 2.5 mm reduced scaffold volume fraction from 41.66% to 14.15%, while porosity increased from 58.34% to 85.85%. Similarly, scaffold strut spacing increased from 0.65 mm to 1.56 mm, qualitatively resembling trabecular deterioration observed in osteoporotic bone [29,38]. Importantly, scaffold volume fraction and scaffold strut spacing represent the corresponding architectural analogs of clinically reported BV/TV and Tb.Sp parameters, respectively.

Benchmarking against HR-pQCT-derived human trabecular bone datasets demonstrated that the intermediate 1.75 mm point spacing scaffold exhibited scaffold volume fraction (24.88%) and porosity (75.12%) values within the clinically reported postmenopausal trabecular bone microarchitectural range [29]. More specifically, the 1.75 mm scaffold aligned near the overlap region between fracture and non-fracture cohorts, whereas the 2.5 mm scaffold represented a more severely deteriorated structural condition with excessive porosity and markedly reduced scaffold volume fraction beyond the average fragility fracture range.

This benchmarking framework is important because it positions the engineered scaffold platform within clinically relevant degenerative architectural states rather than relying solely on arbitrary porosity matching commonly used in scaffold design studies. Prior studies using Voronoi and other lattice-based scaffold designs have largely focused on reproducing trabecular-like morphology, optimizing porosity and mechanical performance, or generating bone-mimetic scaffolds resembling healthy trabecular bone [39,40]. In contrast, the present study establishes Voronoi point spacing as a controllable architectural parameter for generating dense, clinically compromised, and severely deteriorated trabecular-like microenvironments. Thus, the key advance is not simply fabrication of a trabecular-like scaffold, but the use of Voronoi point spacing as a tunable design variable to model architecture-driven mechanical compromise and degeneration-like bone conditions.

Importantly, this study does not claim direct replication of native osteoporotic bone biology. Rather, it establishes an architecturally relevant and quantitatively benchmarked in vitro platform for investigating architecture-dependent biological responses in compromised bone-like microenvironments. Although scaffold parameters were quantified using μCT and compared with clinically reported HR-pQCT datasets, the intent was not to establish imaging equivalence between modalities, but rather to determine whether the engineered scaffold architectures fall within clinically observed trabecular microarchitectural ranges associated with osteoporotic deterioration.

### 4.2 Architecture–cytoskeleton–mineralization coupling and surface-mediated modulation of osteogenic response

The combined confocal imaging and quantitative analyses indicate a consistent relationship between scaffold architecture and early cytoskeletal organization. Lower point spacing conditions (1.25 and 1.75 mm), which exhibit higher structural connectivity and more distributed load transfer, supported greater cell spreading and the formation of more developed actin networks. In contrast, the high point spacing condition (2.5 mm), characterized by reduced connectivity and larger pore gaps, was associated with more limited cell attachment and simplified cytoskeletal organization. The corresponding reductions in actin intensity and stress fiber density suggest that architectural differences are reflected at the level of cytoskeletal organization, consistent with prior observations that cell spreading and actin assembly are influenced by the availability of adhesion sites and mechanical support within 3D environments [30–32].

Previous studies involving Voronoi and other architected lattice scaffolds have primarily focused on reproducing trabecular heterogeneity, optimizing permeability and stiffness, or enhancing osteogenesis within structurally favorable scaffold systems. Similarly, TPMS and gyroid architectures have been widely explored because their interconnected porous geometries support efficient nutrient transport and mechanical stability in bone tissue engineering applications. However, in many of these systems, architectural variation is often accompanied by simultaneous changes in material composition, surface chemistry, or processing conditions, making it difficult to isolate the independent contribution of scaffold architecture to biological response. In contrast, the present framework treats Voronoi point spacing as a single controllable architectural variable within a constant material and fabrication system, enabling systematic investigation of how progressive architectural deterioration alone influences cytoskeletal organization and osteogenic function.

These architectural differences are further supported by μCT-derived structural characterization (Figure 4a–g), which demonstrated a progressive transition toward more porous and less connected architectures with increasing point spacing. Quantitative analysis showed increases in scaffold strut spacing (Tb.Sp-equivalent) and porosity accompanied by decreases in scaffold volume fraction (BV/TV-equivalent), indicating a reduction in structural density and connectivity. Importantly, scaffold volume fraction and scaffold strut spacing represent the corresponding architectural analogs of clinically reported bone morphometry parameters BV/TV and Tb.Sp, respectively. These changes provide a structural basis for the observed differences in cytoskeletal organization, as reduced connectivity likely limits available attachment sites and mechanical continuity within the scaffold network [33,34].

To contextualize these architectural variations, scaffold-derived parameters were benchmarked against clinically reported HR-pQCT datasets from postmenopausal women with and without fragility fractures. The intermediate 1.75 mm condition exhibited scaffold volume fraction and porosity values within the clinically reported postmenopausal trabecular bone microarchitectural range, whereas the 2.5 mm condition extended toward a severely deteriorated structural state characterized by excessive porosity and markedly reduced scaffold volume fraction. Unlike previous scaffold studies that primarily mimic healthy trabecular morphology, the present work establishes a clinically contextualized degeneration-inspired architectural framework spanning dense, compromised, and severely deteriorated trabecular-like microenvironments. This distinction is important because osteoporosis and skeletal fragility are associated not only with reduced bone mass, but also with progressive architectural deterioration and impaired structural connectivity that fundamentally alter load transfer and cellular mechanosensing within bone tissue.

Consistent with these structural and cytoskeletal differences, changes in osteogenic mineralization were observed over time. Scaffolds with lower point spacing exhibited higher and more spatially uniform mineral deposition, whereas the high point spacing condition showed consistently reduced mineralization across time points. Together, these observations support a structure–function relationship in which architectural deterioration is associated with impaired osteogenic outcomes, potentially through its influence on cell–material interactions, cytoskeletal organization, and mechanical continuity within the scaffold microenvironment [20,35].

Building on these architecture-associated limitations, surface biointerface engineering was explored as a strategy to modulate cellular responses without altering scaffold geometry. In this study, polydopamine (PDA) was employed as a surface coating due to its reported osteogenic potential [36,37]. Previous studies have primarily utilized PDA to enhance osteogenesis in structurally favorable scaffolds or as a platform for immobilizing bioactive molecules such as peptides and proteins that promote osteogenic differentiation [38–40]. However, its application within architecturally compromised degeneration-inspired scaffold environments has not been systematically investigated. To the best of our knowledge, the potential of PDA coatings to partially restore osteogenic functionality in architecture-limited microenvironments remains largely unexplored.

SEM analysis indicated that PDA coatings could be applied in a concentration-dependent manner, transitioning from discontinuous surface features at lower concentrations to more uniform coverage at higher concentrations [41]. This progressive change in surface morphology provides a tunable biointerface for cell interaction while preserving the underlying scaffold architecture [42,43]. The three-dimensional confocal reconstructions suggest that scaffold architecture influences not only local cell attachment but also the spatial organization of cells within the scaffold microenvironment. Reduced cytoskeletal continuity observed in the 2.5 mm PS control scaffolds may reflect the reduced structural connectivity associated with architecturally compromised scaffold states. The progressive improvement in cellular organization observed following PDA coating further suggests that surface biointerface engineering can partially modulate cell–material interactions and cytoskeletal organization within fixed architectural conditions without altering bulk scaffold geometry. Functionally, PDA-coated scaffolds exhibited increased mineralization relative to the untreated high point spacing condition, indicating that surface modification partially improved osteogenic response within mechanically compromised scaffold environments. Further more, the observed differences in actin organization and stress fiber density suggest that scaffold architecture influences cytoskeletal organization within the engineered microenvironments. Lower point spacing scaffolds, which exhibited greater structural connectivity, supported more developed actin organization, whereas the high point spacing condition was associated with reduced cytoskeletal continuity. The progressive recovery in stress fiber organization observed following PDA coating further suggests that surface biointerface engineering can partially modulate cytoskeletal organization within architecturally compromised scaffold environments without altering the underlying scaffold geometry.

To quantify this effect, the Mineralization Rescue Percentage (MRP) was introduced as a normalized metric representing the fraction of architecture-associated mineralization reduction recovered following treatment. Using this framework, a concentration-dependent increase in MRP was observed with increasing PDA concentration. However, recovery remained incomplete even at the highest PDA concentration (43% at 1.0% PDA), indicating that surface modification alone does not fully offset the functional limitations associated with severe architectural deterioration. These findings suggest that scaffold geometry and interfacial bioactivity contribute in complementary but non-equivalent ways to osteogenic performance.

A key advance of the present framework is the ability to independently decouple architectural and interfacial contributions to scaffold function. In many biomaterial systems, changes in architecture are inherently accompanied by changes in chemistry, stiffness, or processing conditions, making it difficult to determine whether observed biological responses arise primarily from structural or interfacial effects. Here, scaffold geometry defines the mechanically and structurally compromised microenvironment, while surface biointerface engineering selectively modulates cellular interactions within a fixed architectural state. This experimentally controllable degeneration-to-rescue framework therefore provides a platform for systematically investigating how architectural deterioration and biointerface-mediated rescue collectively regulate osteogenic function.

From a broader perspective, this work has implications for understanding how structural degradation influences regenerative behavior in bone. Osteoporosis and fragility fractures are associated with progressive reductions in trabecular connectivity and mechanical competence, both of which impair the ability of bone tissue to support effective load transfer and regeneration. The observed partial restoration of mineralization through PDA-mediated biointerface engineering suggests that interfacial strategies may provide a means to improve osteogenic responses within mechanically compromised environments, particularly in cases where complete structural restoration is not achievable. Beyond bone tissue engineering, the concept of architecture-defined functional states and their modulation through interface engineering may be broadly applicable to other biological and engineered systems in which connectivity and structural organization govern functional behavior. Collectively, this study establishes a clinically contextualized framework in which a single architectural parameter defines a continuum of degeneration-inspired structural and functional states, while surface biointerface engineering enables partial rescue of osteogenic functionality within these compromised microenvironments.

## 5. Conclusions

This study establishes Voronoi point spacing as a single tunable architectural parameter capable of generating mechanically and structurally distinct scaffold states that model transitions from dense trabecular-like networks to architecturally compromised microenvironments. Increasing point spacing progressively reduced scaffold stiffness, promoted localized stress concentration, disrupted cytoskeletal organization, and suppressed osteogenic mineralization. Benchmarking against clinically reported HR-pQCT datasets further demonstrated the physiological relevance of the engineered architectural states. Importantly, PDA-mediated surface biointerface engineering partially restored cytoskeletal organization and mineralization within structurally compromised scaffolds without altering the underlying architecture. The introduction of the Mineralization Rescue Percentage (MRP) framework further enabled quantitative assessment of biointerface-mediated functional recovery within degeneration-inspired scaffold microenvironments. Collectively, this work provides a clinically contextualized in vitro platform for decoupling architectural and interfacial contributions to osteogenic function and introduces a framework for evaluating bioactive surface interventions under degeneration-relevant structural conditions. These findings establish design principles for architected biomaterials and provide a foundation for future mechanobiological studies and degeneration-relevant biomaterials screening platforms.

## 6. Acknowledgment

This research was supported by the National Science Foundation through the NSF RUI grant (Award No. 2332041) and the Centre for Engineering Mechanobiology (CEMB, CMMI Award No. 15-48571), as well as by the National Institutes of Health through the University of Pennsylvania Institutional Research and Academic Career Development Award (IRACDA, K12GM081259), the Penn Center for Musculoskeletal Disorders (P30-AR069619), and the NIH R01 grant (1R01HD113596) from the National Institute of Arthritis and Musculoskeletal and Skin Diseases. The authors thank Dr. Robert Grabski (Research Scientist, University of Alabama at Birmingham) for his valuable assistance with confocal fluorescence microscopy imaging. The authors thank Dr. Emmanuel Tadjuidje (Professor Biology Department, Alabama State University) for the stereomicroscopic images. V.V. also gratefully acknowledges nTopology for providing an educational software license to Alabama State University.

## 8. Data availability statement

The data that support the findings of this study are available from the corresponding author upon reasonable request.

## References

[1] Y.M. Sillmann, A.M.P. Baggio, P. Eber, B.R. Freedman, C. Liu, Y. Jounaidi, A. Schramm, F. Wilde, F.P.S. Guastaldi, Advancing Scaffold Architecture for Bone Tissue Engineering: A Comparative Study of 3D-Printed β-TCP Constructs in Dynamic Culture with pBMSC, J Funct Biomater 16(9) (2025).

[2] M. Selim, H.M. Mousa, G.T. Abdel-Jaber, A. Barhoum, A. Abdal-hay, Innovative designs of 3D scaffolds for bone tissue regeneration: Understanding principles and addressing challenges, European Polymer Journal 215 (2024) 113251.

[3] T.O. Josephson, E.F. Morgan, Mechanobiological optimization of scaffolds for bone tissue engineering, Biomechanics and Modeling in Mechanobiology 23(6) (2024) 2025–2042.

[4] M.A. Velasco, C.A. Narváez-Tovar, D.A. Garzón-Alvarado, Design, materials, and mechanobiology of biodegradable scaffolds for bone tissue engineering, Biomed Res Int 2015 (2015) 729076.

[5] G. Percoco, A.E. Uva, M. Fiorentino, M. Gattullo, V.M. Manghisi, A. Boccaccio, Mechanobiological Approach to Design and Optimize Bone Tissue Scaffolds 3D Printed with Fused Deposition Modeling: A Feasibility Study, Materials 13(3) (2020) 648.

[6] S. Polo, A. García-Domínguez, E.M. Rubio, J. Claver, Lattice Structures in Additive Manufacturing for Biomedical Applications: A Systematic Review, Polymers (Basel) 17(17) (2025).

[7] G. Reinke, A.C. dos Santos, A systematic review and unified framework for design for additive manufacturing (DfAM), The International Journal of Advanced Manufacturing Technology 143(9) (2026) 4507–4535.

[8] K. Chen, Y. Li, Y. Xuan, M. Khan, X. Wang, X. Zhang, F. Guo, Data-driven multiscale design of composite biomaterials: Integrating experiments, imaging, and computational modeling for biomedical engineering, Materials Today Bio 37 (2026) 102905.

[9] K. Hayashi, T. Yanagisawa, R. Kishida, K. Ishikawa, Effects of Scaffold Shape on Bone Regeneration: Tiny Shape Differences Affect the Entire System, ACS Nano 16(8) (2022) 11755–11768.

[10] M.J. Dewey, R.S.H. Chang, A.V. Nosatov, K. Janssen, S.J. Crotts, S.J. Hollister, B.A.C. Harley, Generative design approach to combine architected Voronoi foams with porous collagen scaffolds to create a tunable composite biomaterial, Acta Biomater 172 (2023) 249–259.

[11] L. D’Andrea, G. Goretti, G. Magrini, P. Vena, Tuning the trabecular orientation of Voronoi-based scaffold to optimize the micro-environment for bone healing, Biomech Model Mechanobiol 24(3) (2025) 1057–1071.

[12] M. Rezapourian, I. Hussainova, Optimal mechanical properties of Hydroxyapatite gradient Voronoi porous scaffolds for bone applications — A numerical study, Journal of the Mechanical Behavior of Biomedical Materials 148 (2023) 106232.

[13] A. Alhelal, D. Ali, M. Hasan, Design and Fabrication of Bone Scaffolds With Regular and Irregular Voronoi Architectures: A Comparative Study, Advances in Polymer Technology 2025(1) (2025) 2529277.

[14] S. Zou, H. Gong, J. Gao, Additively Manufactured Multilevel Voronoi-Lattice Scaffolds with Bonelike Mechanical Properties, ACS Biomaterials Science & Engineering 8(7) (2022) 3022–3037.

[15] L. Chao, Y. He, J. Gu, D. Xie, Y. Yang, L. Shen, G. Wu, L. Wang, Z. Tian, H. Liang, Design of porous structure based on the Voronoi diagram and stress line for better stress shielding relief and permeability, Journal of Materials Research and Technology 25 (2023) 1719–1734.

[16] X. Zhang, M. Zhang, C. Zhang, T. Zhou, X. Wu, X. Yue, Prediction and Numerical Study of Thermal Performance of Gradient Porous Structures Based on Voronoi Tessellation Design, Materials (Basel) 15(22) (2022).

[17] J.G. Garrison, C.L. Slaboch, G.L. Niebur, Density and architecture have greater effects on the toughness of trabecular bone than damage, Bone 44(5) (2009) 924–9.

[18] M.L. Gatto, G. Cerqueni, M. Furlani, N. Riberti, E. Tognoli, L. Denti, F. Leonardi, A. Giuliani, M. Mattioli-Belmonte, P. Mengucci, Influence of Trabecular Geometry on Scaffold Mechanical Behavior and MG-63 Cell Viability, Materials (Basel) 16(6) (2023).

[19] Y. Zhou, P. Isaksson, C. Persson, An improved trabecular bone model based on Voronoi tessellation, Journal of the Mechanical Behavior of Biomedical Materials 148 (2023) 106172.

[20] S.-Y. Long, Y.-J. Fu, Z.-M. Zhang, R. Tang, P. Yu, W. Yang, Architecture mechanics mediated osteogenic progression in bone regeneration of artificial scaffolds, Science Advances 11(29) (2025) eadv8804.

[21] A. Ghasem-Zadeh, M. Bui, E. Seeman, S.K. Boyd, S. Iuliano, R. Jaipurwala, P.F. Mount, N.D. Toussaint, C. Chiang, Bone microarchitecture and estimated failure load are deteriorated whether patients with chronic kidney disease have normal bone mineral density, osteopenia or osteoporosis, Bone 154 (2022) 116260.

[22] T.M. Koushik, C.M. Miller, E. Antunes, Bone Tissue Engineering Scaffolds: Function of Multi-Material Hierarchically Structured Scaffolds, Adv Healthc Mater 12(9) (2023) e2202766.

[23] R. Teimouri, K. Abnous, S.M. Taghdisi, M. Ramezani, M. Alibolandi, Surface modifications of scaffolds for bone regeneration, Journal of Materials Research and Technology 24 (2023) 7938–7973.

[24] G. Osterhoff, E.F. Morgan, S.J. Shefelbine, L. Karim, L.M. McNamara, P. Augat, Bone mechanical properties and changes with osteoporosis, Injury 47 Suppl 2(Suppl 2) (2016) S11–20.

[25] B.M. Mulvihill, L.M. McNamara, P.J. Prendergast, Loss of trabeculae by mechano-biological means may explain rapid bone loss in osteoporosis, J R Soc Interface 5(27) (2008) 1243–53.

[26] J. Wang, Y. Cui, B. Zhang, S. Sun, H. Xu, M. Yao, D. Wu, Y. Wang, Polydopamine-Modified functional materials promote bone regeneration, Materials & Design 238 (2024) 112655.

[27] T.S. Sileika, H.-D. Kim, P. Maniak, P.B. Messersmith, Antibacterial Performance of Polydopamine-Modified Polymer Surfaces Containing Passive and Active Components, ACS Applied Materials & Interfaces 3(12) (2011) 4602–4610.

[28] M. Huo, S. He, Y. Zhang, Y. Feng, J. Lu, Simulation on bone remodeling with stochastic nature of adult and elderly using topology optimization algorithm, Journal of Biomechanics 136 (2022) 111078.

[29] X.S. Liu, E.M. Stein, B. Zhou, C.A. Zhang, T.L. Nickolas, A. Cohen, V. Thomas, D.J. McMahon, F. Cosman, J. Nieves, E. Shane, X.E. Guo, Individual trabecula segmentation (ITS)-based morphological analyses and microfinite element analysis of HR-pQCT images discriminate postmenopausal fragility fractures independent of DXA measurements, Journal of Bone and Mineral Research 27(2) (2012) 263–272.

[30] S. Kanwar, O. Al-Ketan, S. Vijayavenkataraman, A novel method to design biomimetic, 3D printable stochastic scaffolds with controlled porosity for bone tissue engineering, Materials & Design 220 (2022) 110857.

[31] J. Li, D. Guo, J. Li, X. Wei, Z. Sun, B. Yang, T. Lu, P. Ouyang, S. Chang, W. Liu, X. He, Irregular pore size of degradable bioceramic Voronoi scaffolds prepared by stereolithography: Osteogenesis and computational fluid dynamics analysis, Materials & Design 224 (2022) 111414.

[32] L. D’Andrea, M. Montemurro, P. Castany, Tuning the trabecular orientation of Voronoi-based scaffold to mimic bone anisotropy for tissue engineering applications, Biomechanics and Modeling in Mechanobiology 24 (2025) 1–17.

[33] M. Shen, Y. Li, F. Lu, Y. Gou, C. Zhong, S. He, C. Zhao, G. Yang, L. Zhang, X. Yang, Z. Gou, S. Xu, Bioceramic scaffolds with triply periodic minimal surface architectures guide early-stage bone regeneration, Bioactive Materials 25 (2023) 374–386.

[34] A. Yánez, A. Cuadrado, O. Martel, M.P. Fiorucci, S. Deviaene, Mechanical and permeability properties of skeletal and sheet triply periodic minimal surface scaffolds in bone defect reconstruction, Results in Engineering 21 (2024) 101883.

[35] J. Ma, Y. Li, Y. Mi, Q. Gong, P. Zhang, B. Meng, J. Wang, J. Wang, Y. Fan, Novel 3D printed TPMS scaffolds: Microstructure, characteristics and applications in bone regeneration, Journal of Tissue Engineering 15 (2024) 20417314241263689.

[30] P.T. Caswell, T. Zech, Actin-Based Cell Protrusion in a 3D Matrix, Trends Cell Biol 28(10) (2018) 823–834.

[31] X. Wang, Y. Yang, Y. Wang, C. Lu, X. Hu, N. Kawazoe, Y. Yang, G. Chen, Focal adhesion and actin orientation regulated by cellular geometry determine stem cell differentiation via mechanotransduction, Acta Biomaterialia 182 (2024) 81–92.

[32] A.D. Doyle, K.M. Yamada, Mechanosensing via cell-matrix adhesions in 3D microenvironments, Exp Cell Res 343(1) (2016) 60–66.

[33] V.D.L. Putra, K.A. Kilian, M.L. Knothe Tate, Biomechanical, biophysical and biochemical modulators of cytoskeletal remodelling and emergent stem cell lineage commitment, Communications Biology 6(1) (2023) 75.

[34] N. Chen, M. Wu, R. Williams, J. Yan, J. Zhou, D. Zhu, Y. Ding, Strong living scaffolds for load-bearing musculoskeletal tissue regeneration, Mater Today Bio 35 (2025) 102571.

[35] G. Ratheesh, M. Shi, P. Lau, Y. Xiao, C. Vaquette, Effect of Dual Pore Size Architecture on In Vitro Osteogenic Differentiation in Additively Manufactured Hierarchical Scaffolds, ACS Biomaterials Science & Engineering 7(6) (2021) 2615–2626.

[36] S. Boutroy, M.L. Bouxsein, F. Munoz, P.D. Delmas, In vivo assessment of trabecular bone microarchitecture by high-resolution peripheral quantitative computed tomography, Journal of Clinical Endocrinology and Metabolism 90 (2005) 6508–6515.

[37] E. Sornay-Rendu, S. Boutroy, F. Munoz, P.D. Delmas, Alterations of cortical and trabecular architecture are associated with fractures in postmenopausal women, independent of bone mineral density, Journal of Bone and Mineral Research 22 (2007) 425–433.

[38] N. Vilayphiou, S. Boutroy, P. Szulc, B. Van Rietbergen, F. Munoz, P.D. Delmas, R. Chapurlat, Finite element analysis performed on HR-pQCT images of bone microarchitecture in postmenopausal women is associated with distal radius fracture status, Journal of Bone and Mineral Research 25 (2010) 983–990.

[39] M. Fantini, M. Curto, F. De Crescenzio, A method to design biomimetic scaffolds for bone tissue engineering based on Voronoi lattices, Virtual and Physical Prototyping 11 (2016) 77–90.

[40] B. Herath, T. Downing, S. Yin, R. Li, X. Wang, Mechanical and geometrical study of 3D printed Voronoi scaffold design for large bone defects, Materials & Design 212 (2021) 110224.

[41] M.L. Alfieri, L. Panzella, S.L. Oscurato, M. Salvatore, R. Avolio, M.E. Errico, P. Maddalena, A. Napolitano, M. D’Ischia, The Chemistry of Polydopamine Film Formation: The Amine-Quinone Interplay, Biomimetics (Basel) 3(3) (2018).

[42] W.H. Song, J.E. Kim, L. Rajbongshi, S.R. Lee, Y. Kim, S.Y. Hwang, S.O. Oh, B.S. Kim, D. Lee, S. Yoon, Polydopamine-Coated Surfaces Promote Adhesion, Migration, Proliferation, Chemoresistance, Stemness, and Epithelial-Mesenchymal Transition of Human Prostate Cancer Cell Lines In Vitro via Integrin α(2)β(1)-FAK-JNK Signaling, Int J Mol Sci 27(2) (2026).

[43] Z. Deng, W. Wang, X. Xu, Y. Nie, Y. Liu, O.E.C. Gould, N. Ma, A. Lendlein, Biofunction of Polydopamine Coating in Stem Cell Culture, ACS Applied Materials & Interfaces 13(9) (2021) 10748–10759.

