## Supporting Information for "Degeneration-Inspired Architectural States Defined by Voronoi Point Spacing and Surface-Mediated Rescue of Osteogenic Dysfunction in 3D-Printed Scaffolds"

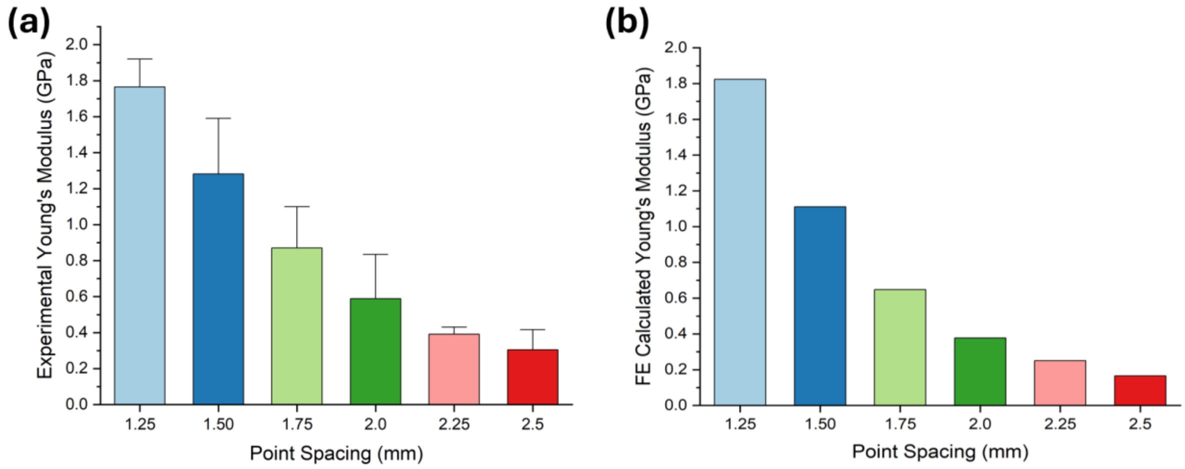


**Figure S1:** Experimental and finite element (FE)-predicted Young's modulus of Voronoi scaffolds with varying point spacing (1.25–2.5 mm). (a) Experimental Young's modulus obtained from compression testing. (b) FE-calculated Young's modulus. Both analyses demonstrate progressive reduction in scaffold stiffness with increasing point spacing.

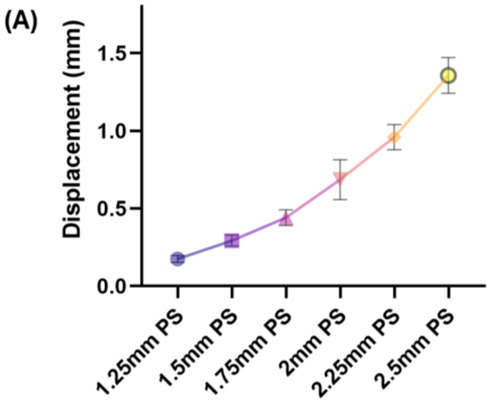


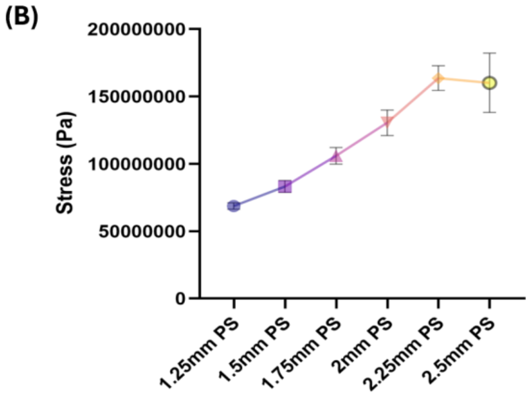


(C)

| Point Spacing (mm) | Mean Stress (Pa) | Std Dev (Pa) | Max Stress (Pa) | Max/Mean Stress Ratio |
|---|---|---|---|---|
| 1.25 PS | 6.87E+07 | 6.18E+06 | 6.36E+08 | 9.26 |
| 1.5 PS | 8.32E+07 | 1.20E+07 | 1.09E+09 | 13.10 |
| 1.75 PS | 1.06E+08 | 1.64E+07 | 1.38E+09 | 13.02 |
| 2 PS | 1.31E+08 | 2.49E+07 | 1.85E+09 | 14.12 |
| 2.25 PS | 1.64E+08 | 2.42E+07 | 2.26E+09 | 13.78 |
| 2.5 PS | 1.60E+08 | 5.83E+07 | 3.13E+09 | 19.56 |

**Figure S2.** Quantitative analysis of finite element modeling (FEM) results for Voronoi scaffolds with varying point spacing (1.25–2.5 mm). (A) Mean displacement (mm) as a function of point spacing, showing an increase in deformation with increasing architectural spacing. (B) Mean von Mises stress (Pa) across point spacing conditions, with error bars representing standard deviation. (C) Summary of stress distribution metrics, including mean stress, standard deviation, maximum stress, and maximum-to-mean stress ratio, highlighting increased stress variability and concentration at higher point spacings.

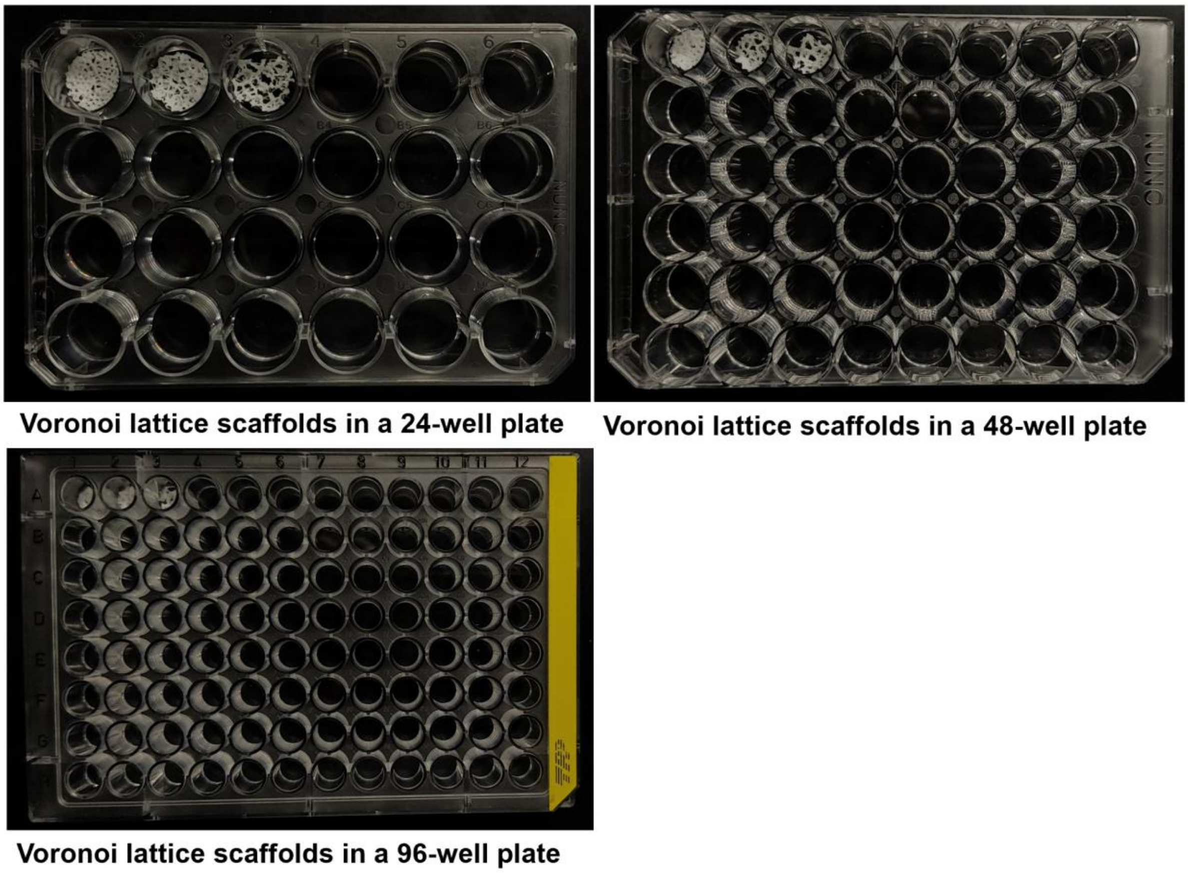


**Figure S3.** Representative placement of 3D-printed Voronoi lattice scaffolds within standard multiwell culture formats. Voronoi scaffolds are shown positioned in (A) 24-well, (B) 48-well, and (C) 96-well plates, demonstrating compatibility with commonly used in vitro platforms and scalability for higher-throughput experimental configurations.

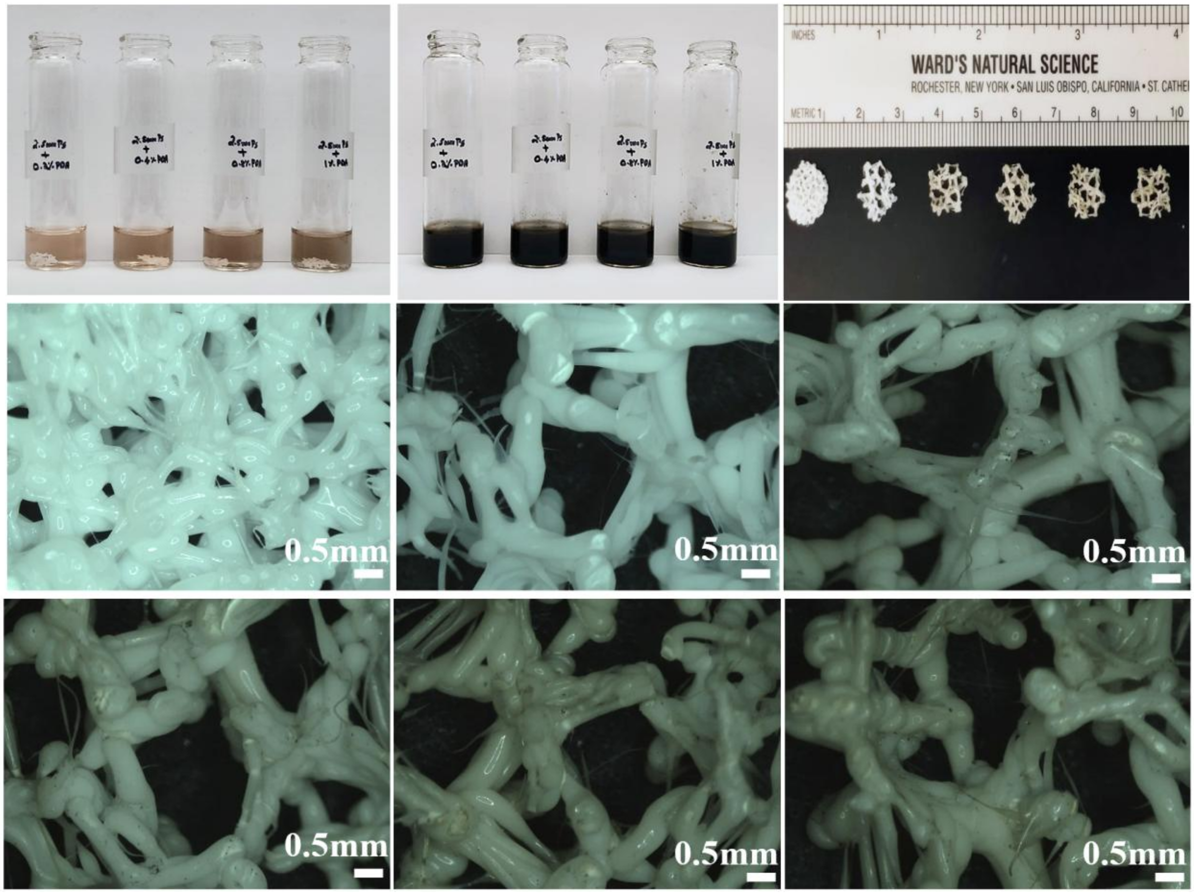


**Figure S4.** PDA coating and morphology of Voronoi scaffolds. Top row shows 2.5 mm scaffolds incubated in dopamine hydrochloride solutions (0.2%, 0.4%, 0.8%, and 1.0% PDA, left to right), exhibiting progressive darkening after 18 h. Top right: macroscopic scaffold views across point spacings. Middle and bottom rows (scale bar: 0.5 mm) show stereomicroscopic images of 1.25 mm control, uncoated 2.5 mm scaffolds, and PDA-coated 2.5 mm scaffolds, highlighting concentration-dependent surface modification with preserved architecture.
